# Thyroid hormones maintain parvalbumin neuron functions in the mouse neocortex

**DOI:** 10.1101/2024.07.16.603713

**Authors:** Juan Ren, Suzy Markossian, Romain Guyot, Denise Aubert, Fabrice Riet, Jacques Brocard, Alice Descamps, Jiemin Wong, Dongdong Li, Bruno Cauli, Thierry Gallopin, Frédéric Flamant, Sabine Richard

## Abstract

Parvalbumin-expressing GABAergic interneurons play a key role in maintaining the excitation-inhibition balance in the mammalian neocortex. While postnatal maturation of parvalbumin neurons is highly sensitive to thyroid hormone deficiency, the role of these hormones in mature parvalbumin neurons remains poorly understood. Here, we addressed the possibility that thyroid hormones are also required to maintain the function of mature parvalbumin neurons in the mouse neocortex. To this end, Cre/loxP recombination system was used to express a dominant-negative mutated receptor of thyroid hormones to selectively eliminate thyroid hormone signaling in parvalbumin neurons after the onset of parvalbumin expression. We analyzed the neocortical phenotype of these mice by combining genomics, histology, electrophysiology, sleep recordings and behavioral analysis. In the mutant mice, gene expression analysis revealed a decreased expression of key perineuronal net components. parvalbumin neuron excitability was decreased and mice displayed behavioral hyperactivity and increased susceptibility to epileptic seizures. Additionally, sleep analysis revealed that mutant mice spent more time awake, and had a delayed transition to sleep. Electrophysiological recordings further demonstrated altered gamma oscillations in the somatosensory cortex. The findings of this study underscore that thyroid hormones are essential not only for the differentiation of parvalbumin interneurons, but also for the maintenance of their inhibitory function after the onset of parvalbumin expression. As a consequence, alterations in thyroid hormone signaling, during development or in adulthood, may contribute to the occurrence of neurological and psychiatric disorders involving altered cortical oscillations.

## Introduction

The dynamic balance between excitatory and inhibitory neurotransmissions plays a fundamental role in the function of neocortical neuronal circuits in mammals (1). Inhibitory GABAergic interneurons account for approximately 15% of the neuronal population of the neocortex (2). They control the excitatory signaling of neighboring pyramidal neurons and prevent runaway feed-forward excitation. Among neocortical GABAergic interneurons, parvalbumin-expressing (PV) neurons have a fast-spiking phenotype (3). They provide powerful inhibition to excitatory neurons, which is crucial for generating gamma oscillations (30–100 Hz) in the neocortex (4). By synchronizing the activity of pyramidal neurons, they facilitate rhythmic patterns that support a variety of cognitive processes. Alteration of their function notably alters the homeostatic regulation of sleep (5) and results in anxiety-like behavior (6).

The appearance of parvalbumin in the neocortex, which occurs during the second postnatal week in mice (7), is a relatively late event of neocortical circuit maturation. It is highly dependent on the presence of thyroid hormones (THs) (8). THs (thyroxine, or T4, and 3,3’,5-triiodo-L-thyronine, or T3, its active metabolite) mainly act by binding to nuclear receptors called TRα1, TRβ1, and TRβ2, which are encoded by the *Thra* and *Thrb* genes in mice. These receptors, collectively called TRs, act as TH-dependent transcription factors, activating the expression of target genes. In both humans and mice, *Thra* mutations have dramatic consequences on development, notably neurodevelopment (9–11). In the neocortex, the most visible consequence of *Thra* mutations is an impairment of PV neuron maturation, as evidenced by a strong reduction in the expression of parvalbumin (12, 13).

Currently, it is unclear whether the requirement for TH signaling persists in PV neurons once their maturation is complete. The alternative would be that THs are necessary for PV neurons only during a short postnatal period, and that early alterations would have long-lasting consequences. We address here this question by using a mouse model in which a mutated *Thra* allele, *Thra^AMI^* (14), drives the expression of the TRα1^L400R^ dominant-negative receptor exclusively in PV neurons (*Thra^AMI/+^ Pvalb-Cre* mice). This leads to a specific blockade of TH signaling in PV neurons after the onset of *Pvalb* gene expression, which encodes parvalbumin. Unlike mice that express this mutation at an earlier stage in all GABAergic neurons (13), *Thra^AMI/+^ Pvalb-Cre* mice were viable and did not display any major neurological defect. Multiscale phenotyping of *Thra^AMI/+^ Pvalb-Cre* mice, which combined genomic, histological, electrophysiological, and behavioral observations, revealed alterations in the functions of neocortical PV neurons. These observations demonstrate that TH signaling in PV neurons is required for their proper function in the adult neocortex.

## Results

### Genomic consequences of TRα1^L400R^ expression in PV neurons

Our first aim was to analyze the long-term and cell-autonomous consequences of the inhibition of TH/TRα1 signaling in PV neurons, using gene expression analysis. Therefore, we combined the floxed *Thra^AMI^* and *Pvalb-Cre* transgenes with the *ROSA-GS-GFP-L10^lox^* reporter construct (Supplementary Fig. 1), which produces a green fluorescent protein in the nuclei of PV neurons after Cre/loxP recombination. Fluorescent cell nuclei were sorted from the neocortex of adult mice expressing the dominant-negative TRα1^L400R^ in PV neurons and from the cortex of control littermates (*Thra^AMI^*^/*+*^ *ROSA-GS-GFP-L10^lox/+^ Pvalb-Cre* mice vs *Thra^+^*^/*+*^ *ROSA-GS-GFP-L10^lox/+^ Pvalb-Cre* control littermates). RNA was extracted from the sorted nuclei and used for RNA-Seq. Differential gene expression analysis identified 281 genes whose expression was either increased or decreased in PV neurons of *Thra^AMI^*^/*+*^ *ROSA-GS-GFP-L10^lox/+^ Pvalb-Cre* mice (Fig. 1A).

**Fig. 1.**
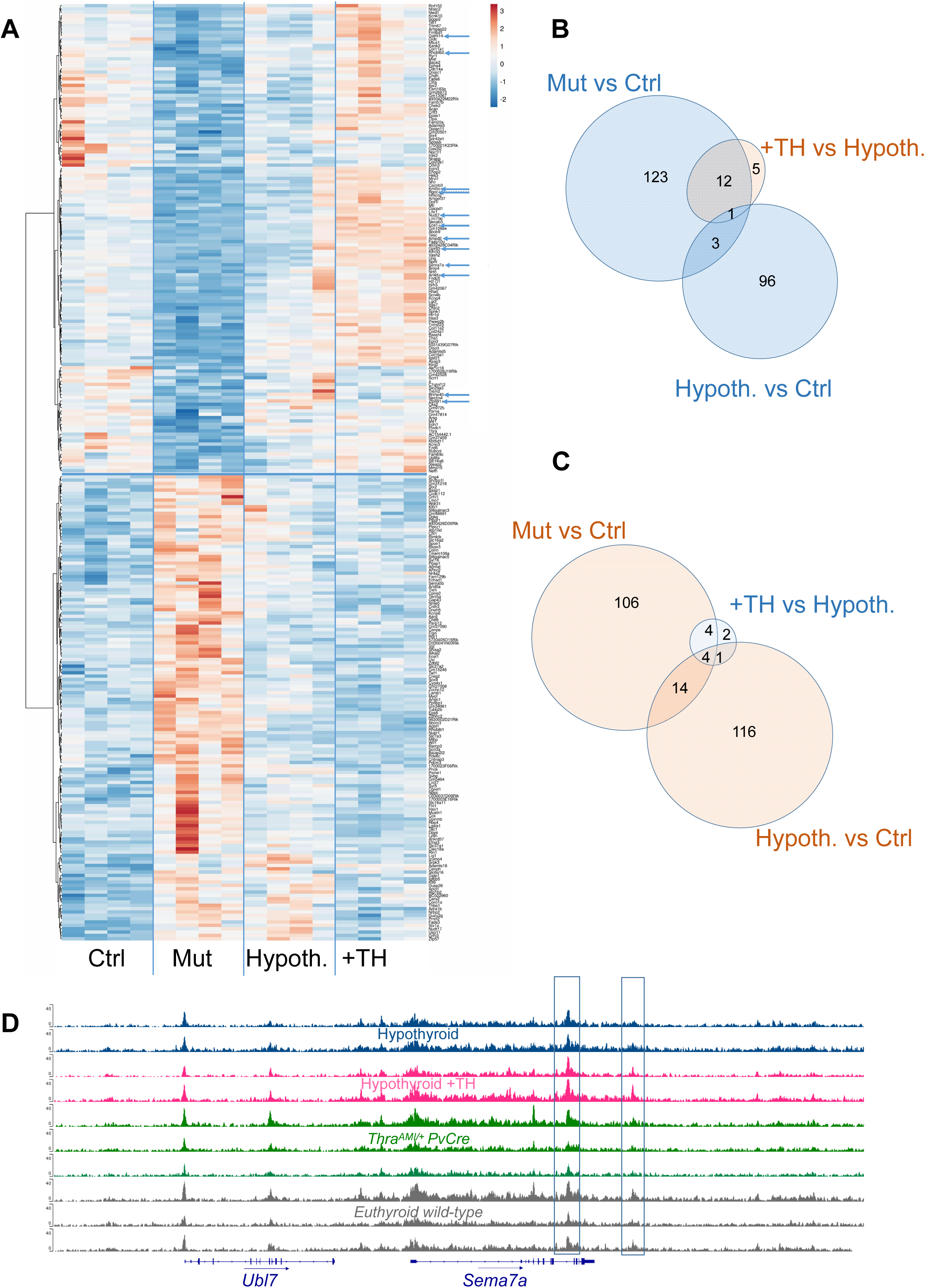
Changes in gene expression and chromatin condensation caused by TRα1^L400R^ expression in PV neurons. **A.** RNA-Seq analysis of nuclear transcriptome of PV neurons. The heatmap represents the variation in gene expression (blue: downregulation, orange: upregulation) of the 281 genes that are differentially expressed in *Thra^AMI^*^/*+*^ *ROSA-GS-GFP-L10^lox^ Pvalb-Cre* mice (Mut, n = 4) and *Thra^+^*^/*+*^ *ROSA-GS-GFP-L10^lox^ Pvalb-Cre* control littermates (Ctrl, n = 4). Although less visible, hypothyroidism has a similar influence on their expression level (Hypoth., n = 4) to that of TRα1^L400R^ expression. TH treatment tends to restore the original level of gene expression, although only partially after 24 hours (+TH; n = 4). Blue arrows indicate the 12 genes for which ATAC-seq indicates a modification of chromatin accessibility caused by the binding of TH to TRs in a sequence located within 30 kb of the transcription start site (see text). **B.** Overlap between the changed responses to TRα1^L400R^ expression (Mut vs Ctrl), hypothyroidism (Hypoth. vs Ctrl) and TH treatment of hypothyroid mice (+TH vs Hypoth.) in the above RNA-Seq analysis. The numbers represent the genes that are down-regulated by reduced thyroid signaling and/or up-regulated upon TH treatment. Here we applied a threshold of 1.5 for fold-changes (blue: down-regulated, orange: upregulated). 4 genes which expression does not fit with these categories are not represented. Note that 24 hours of TH treatment is not sufficient for a full restoration of gene expression in hypothyroid mice. **C.** Same analysis for genes that are up-regulated by reduced thyroid signaling and/or down-regulated upon TH treatment. **D.** An example of ATAC-seq results next to the *Sema7a* gene. The likely positions of TR binding sites in PV neurons are boxed. Note the influence of TH treatment on peak height, which is statistically significant for the two boxed peaks. Also see Supplementary Table 1 for related analyses and datasets.

We then attempted to make a distinction among these 281 genes between direct TR target genes, whose transcription is normally up-regulated by the liganded TRs, and genes whose deregulation is only secondary to the long-term expression of TRα1^L400R^. In order to distinguish between these two categories, we searched for genes whose expression is rapidly modified in PV neurons when adult hypothyroid mice are treated with THs. Hypothyroidism was induced in a group of adult *ROSA-GS-GFP-L10^lox/+^ Pvalb-Cre* mice by a prolonged pharmacological treatment. Some of these hypothyroid mice then received a single intraperitoneal injection of THs to restore the circulating levels of THs, and gene expression was analyzed 24 hours later. In this setting, TH response takes place in all cell types and influences gene expression in PV neurons both directly and indirectly, as it modifies their microenvironment. In sorted nuclei of PV neurons, only 9 genes (*Srpk3, Izumo4, Prmt2, Gucy2g, Gm13067, Gpcpd1, Acan, Stac2, Camk1)* behaved like *bona fide* TR target genes in PV neurons, being equally sensitive to hypothyroidism and TRα1^L400R^ expression, while sensitive to TH treatment (Fig. 1B, C and Supplementary Table 1). Of note, the statistical filters that we have used to identify TR target genes are very stringent, so that the transcriptome analysis that we performed probably underestimates the number of TR target genes in PV neurons.

Another approach to pinpoint TR target genes would be to identify the TR binding sites (TRBSs) in chromatin. We could not use a ChIP-Seq analysis to directly analyze chromatin occupancy by TRs, because of the limited amount of chromatin extracted from sorted nuclei of PV neurons. Therefore, we chose an indirect approach, relying on the fact that chromatin opening occurs in a ligand-dependent manner near TRBSs (15, 16). We thus used ATAC-seq to assess the influence of THs on chromatin accessibility in PV neurons (Fig. 1D). Differential data analysis (hypothyroid mice *vs* hypothyroid mice treated with THs) identified 11 207 differentially accessible sites (DASs, that were more accessible after TH treatment), representing ∼25% of all the detected accessible sites. However, the set of genes located near the DASs (< 30 kb), was not enriched in genes that were identified as differentially expressed after TH treatment. Other comparisons (mutant *vs* control mice; hypothyroid *vs* control mice) identified only 54 DASs, and none of them was proximal to a TH-regulated gene. These data indicate that only a fraction of DASs is associated to TRBSs, and that ATAC-seq alone is not sufficient to identify TR target genes.

In order to predict the position of at least some of the TRBSs in PV neurons, we assumed that a fraction of these would be shared across cell types, as indicated by a recent atlas of ChiP-Seq datasets (17). Using Zekri et al.’s ChiP-Seq atlas, we extracted 4 181 TRBSs that are shared by GABAergic neurons and at least one unrelated cell type (cardiomyocyte, hepatocyte, or adipocyte), from the total 18 064 TRBSs present in striatal GABAergic neurons. We then focused on 1 013 of these 4 181 TRBSs that overlap with the above-defined DASs in PV neurons (called TRBS/DAS), and compared them to the 281 genes that we found to be differentially expressed in *Thra^AMI^*^/*+*^ *ROSA-GS-GFP-L10^lox/+^ Pvalb-Cre* mice (see above). This allowed to pinpoint 12 genes (*Galnt14, Rhobtb2, Kmt5c, Rgcc, Nudt7, Ece1, Ampd2, Gpr83, Sema7a*, *Arl4d, Bhlhe40, Zfp691*) that fulfill two criteria indicating that they are TR target genes: they are down-regulated in *Thra^AMI^*^/*+*^ *ROSA-GS-GFP-L10^lox/+^ Pvalb-Cre* mice, and their transcription start site is located within 30 kb of a TRBS/DAS. Importantly, none of the TRBS/DAS was located near a gene up-regulated in TRα1^L400R^ expressing PV neurons or down-regulated by THs, which is consistent with the fact that liganded TRs activate transcription.

The 281 differentially expressed genes that were identified in TRα1^L400R^-expressing PV neurons have various known functions in signal transduction, ion exchanges and enzymatic activities. Among them, 31 genes (11%) encode proteins that are found in the extracellular domain, mainly extracellular matrix components, of which several fulfil several of the criteria used above to recognize TR target genes (*Acan, Npnt, Sema7a, Vash2, Tesc,* see Supplementary Table 1). This suggests a possible defect in the perineuronal net (PNN), a specialized and highly glycosylated extracellular matrix that surrounds many cortical PV neurons. PNNs provide PV neurons with a mesh-like structure accommodating numerous synaptic terminals, a physical barrier between the neuronal surface and the extracellular space, and its diverse interacting molecules. These functions allow PNNs to regulate the plasticity of PV neurons (18).

### Histological consequences of TRα1^L400R^ expression in PV neurons

We have previously found that PV neurons display a marked sensitivity to alterations in TH signaling during brain development (13). We thus set out to study if the density of PV neurons in the adult mouse neocortex differed between *Thra^AMI^*^/*+*^ *ROSA-tdTomato^lox/+^ Pvalb-Cre* mice (n = 9) and *Thra^+^*^/*+*^ *ROSA-tdTomato^lox/+^ Pvalb-Cre* control littermates (n = 7), which produce a red fluorescent protein in the cytoplasm of PV neurons after Cre/loxP recombination. Immunohistochemistry was performed to label parvalbumin and Wisteria floribunda agglutinin (WFA) was used to label N-acetyl-galactosamine-β1 residues of PNN glycoproteins in the somatosensory cortex (SomCx).

We found no difference between the two groups of mice in the density of Tomato-expressing cells (Fig. 2A, B), which indicates that the expression of TRα1^L400R^ did not compromise the viability of the targeted cells, namely PV neurons. Compared to controls, mutant mice exhibited a slightly, but significantly, higher density of parvalbumin-expressing cells (Fig. 2A, C), accompanied by a slightly, but significantly, lower ratio of parvalbumin-expressing cells that were surrounded by PNNs (Fig. 2A, E). Thus, unlike what we have previously observed in mice with TRα1^L400R^ expressed from a prenatal stage in all GABAergic neurons (13), the differentiation of PV neurons is not drastically altered in *Thra^AMI^*^/*+*^ *ROSA-tdTomato^lox/+^ Pvalb-Cre* mice. Since PNNs surrounding PV neurons allow them to regulate plasticity, the above results suggest that TH signaling in PV neurons might be of physiological importance in the regulation of neural plasticity in the neocortex.

**Fig. 2.**
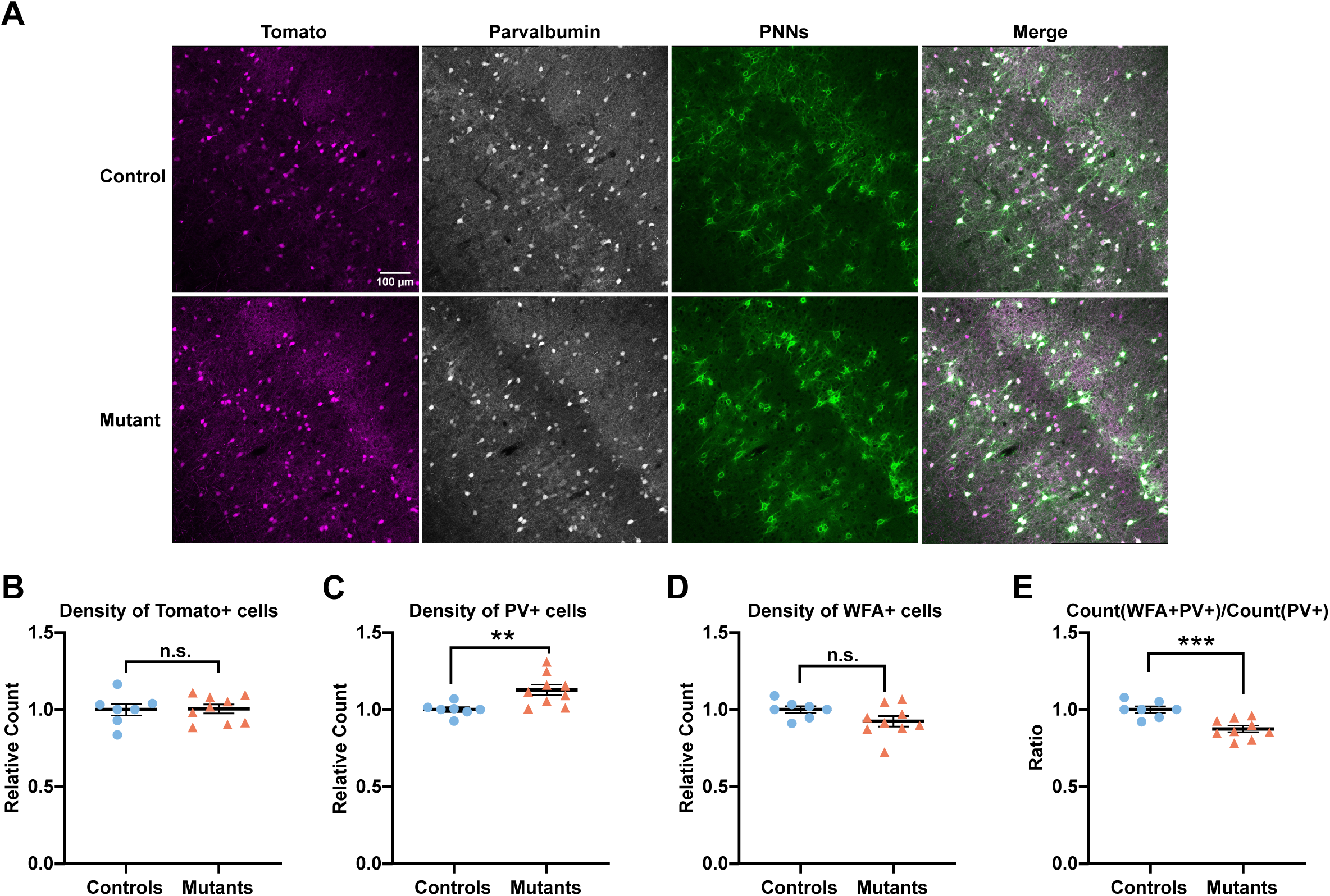
Parvalbumin and PNN alterations in PV neurons expressing TRα1^L400R^ in the somatosensory cortex. **A.** Immunohistochemistry of parvalbumin and WFA-labeled PNNs in PV neurons of somatosensory cortex. **B.** The density of Tomato+ cells in *Thra^AMI^*^/*+*^ *ROSA-tdTomato^lox/+^ Pvalb-Cre* mice (n = 9) did not differ from that in *Thra^+^*^/*+*^ *ROSA-tdTomato^lox/+^ Pvalb-Cre* control littermates (n = 7). **C.** The density of PV+ cells was slightly, but significantly, higher in mutants. **D.** The density of WFA+ cells did not differ between the two groups. **E.** The proportion of PV neurons surrounded by PNNs was slightly, but significantly lower in mutants. Data are presented as mean ± SEM and analyzed using unpaired two-tailed T-test (\*\**p* < 0.01, \*\*\**p* < 0.001, n.s. *p* > 0.05).

Fast-spiking properties of PV cells impose a heavy metabolic burden, rendering them susceptible to oxidative stress, against which PNNs have a protective effect (19). As the percentage of PV neurons surrounded by PNNs were significantly lower in *Thra^AMI^*^/*+*^ *ROSA-tdTomato^lox/+^ Pvalb-Cre* mice than in control mice, we hypothesized that long-term expression of TRα1^L400R^ in PV neurons might increase oxidative stress in these neurons. To test this hypothesis, we labeled cortical sections with an antibody directed against the oxidative stress marker 8-oxo-2’-deoxyguanosine (8-oxo-dG). There was no difference between *Thra^AMI^*^/*+*^ *ROSA-tdTomato^lox/+^ Pvalb-Cre* and control mice in the density of Tomato+ 8-oxo-dG+ cells, nor in the intensity of 8-oxo-DG labelling in Tomato+ cells (Supplementary Fig. 2).

### Whole-cell patch-clamp recordings of PV neurons expressing TRα1^L400R^

The fact that TRα1^L400R^ expression in PV neurons deregulated genes encoding PNN components and ions channels, and reduced the ratio of PV neurons surrounded by PNNs, prompted us to evaluate the electrophysiological properties of PV neurons expressing TRα1^L400R^. Whole-cell current-clamp recording was performed on acute slices of the SomCx, which were prepared from 16-to-24-day-old *Thra^AMI^*^/*+*^ *ROSA-tdTomato^lox/+^ Pvalb-Cre* mice (n = 10) and *Thra^+^*^/*+*^ *ROSA-tdTomato^lox/+^ Pvalb-Cre* control littermates (n = 8). Current steps were injected to fluorescent cells, i.e. PV neurons, and voltage responses were recorded (Fig. 3A, B). In the end, 35 mutant and 19 control PV neurons were recorded, and 37 parameters were measured for each cell (20) in accordance with Petilla terminology (21). This allowed for a comprehensive analysis of the intrinsic electrophysiological behavior of each recorded PV neuron (Supplementary Table 2).

**Fig. 3.**
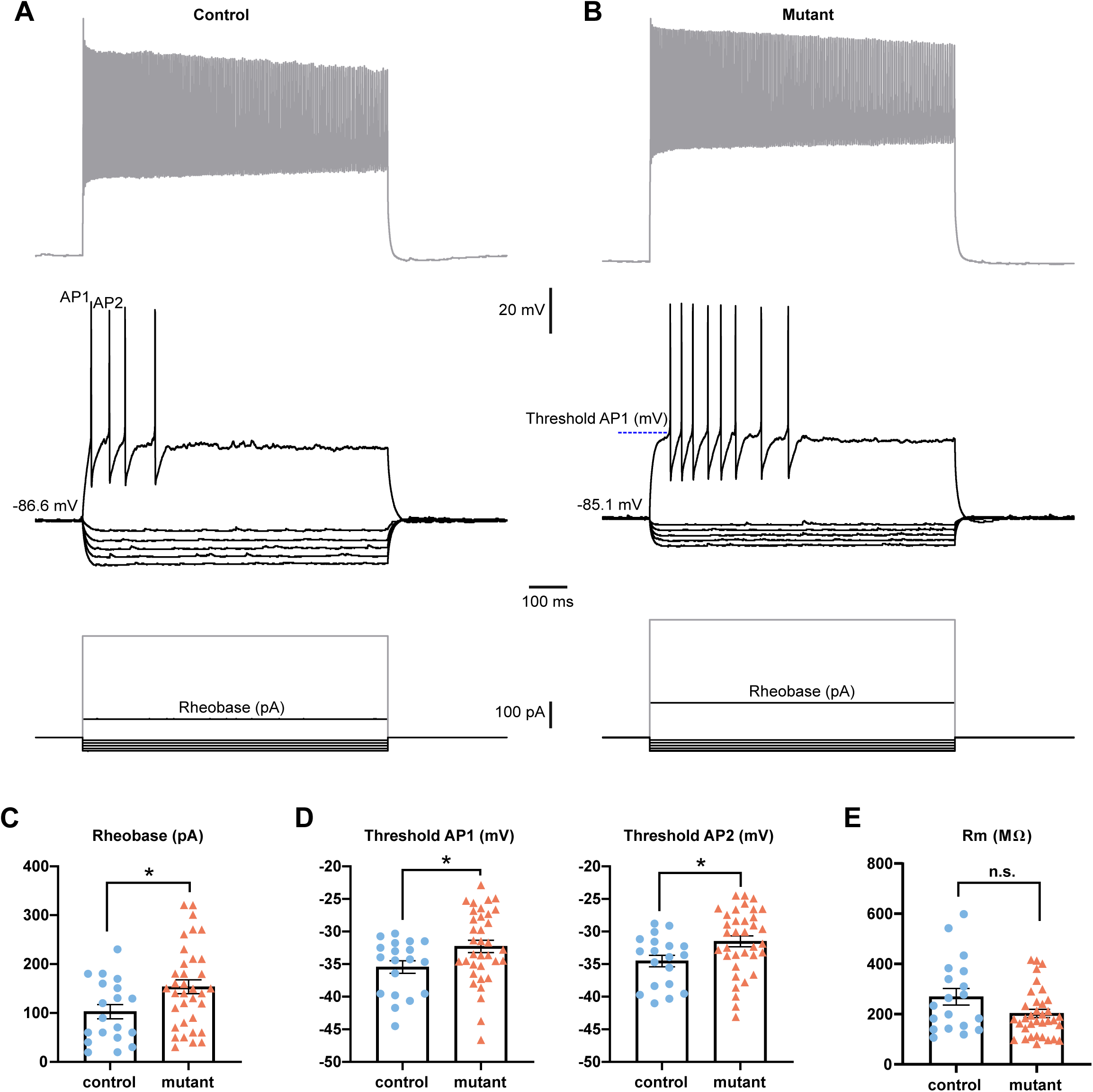
Electrophysiological alterations in PV neurons expressing TRα1^L400R^ in the somatosensory cortex. **A.** Current-clamp recordings of a control (*Thra^+^*^/*+*^ *ROSA-tdTomato^lox/+^ Pvalb-Cre*) PV neuron in response to the current pulses (bottom traces, increased from bottom to top) of -100, -80, - 60, -40, -20, +140, and +760 pA. Note the short latency of fast action potentials (APs) (middle trace) induced by a just-above-threshold current pulse (140 pA). Strong depolarizing current (760 pA) evoked a high and saturated firing rate (top trace). **B.** Current-clamp recordings of a mutant (*Thra^AMI^*^/*+*^ *ROSA-tdTomato^lox/+^ Pvalb-Cre*) PV neuron in response to the current pulses (bottom traces, increased from bottom to top) of -100, -80, -60, -40, -20, +260, and +880 pA. Note the short latency of fast APs (middle trace) induced by a just-above-threshold current pulse (260 pA). Strong depolarizing current (880 pA) evoked a high and saturated firing rate (top trace). **C.** Rheobase, the minimal current intensity required for AP firing, is higher in mutant PV neurons (n = 35) than that in the controls (n = 19). **D.** The thresholds of the first two APs induced by the just-above-threshold current pulse in mutant PV neurons (n = 35), are higher than those in the controls (n = 19). **E** Membrane resistance did not differ significantly between mutant and control PV neurons. Data are presented as mean ± SEM and were analyzed using unpaired two-tailed T-test (\**p* < 0.05, *n.s.* p > 0.1)). Also see Supplementary Table 2 for related analyses and datasets.

Recorded neurons displayed the archetypal properties of fast-spiking PV neurons, with a relatively low membrane resistance (R_m_), short spikes with sharp after-hyperpolarizations (AHPs), and an ability to fire action potentials at a high firing rate with little or no frequency adaptation (Fig. 3A, B and supplementary table 2). Three parameters differed significantly in the PV neurons of *Thra^AMI^*^/*+*^ *ROSA-tdTomato^lox/+^ Pvalb-Cre* mice, compared to controls. The rheobase (Fig. 3C), which is the minimal depolarizing current pulse intensity generating at least one action potential, was significantly higher in mutant PV neurons. The two other parameters that were also higher in mutants were the thresholds of the first and second action potential generated by a rheobasic current injection (Fig. 3D). Membrane resistance did not differ significantly between mutant and control PV neurons (Fig. 3E). Collectively, these alterations indicate that PV neurons expressing TRα1^L400R^ exhibit a lower excitability.

### In vivo local field potential recording of cortical electrical activity

PV neurons play a crucial role in the generation and maintenance of gamma oscillations, as they are key regulators of inhibitory network synchronization. Given their central role in circuit dynamics, we sought to determine whether gamma activity was modified in mutant mice, potentially reflecting impairments in inhibitory network function. To this end, we implanted electrodes to record local field potential (LFP) *in vivo* in adult *Thra^AMI/+^ ROSA-tdTomato^lox/+^ Pvalb-Cre* mutant mice (n = 4) and their *Thra^+/+^ ROSA-tdTomato^lox/+^ Pvalb-Cre* control littermates (n = 4) and assessed spectral activity. In parallel, we characterized and quantified vigilance states in mutant mice to gain insights into potential disruptions in sleep architecture and state-dependent oscillatory dynamics.

To determine whether mutant mice exhibit alterations in sleep-wake regulation, we analyzed the proportion of time spent in wakefulness, NREM sleep, and REM sleep, as well as sleep onset latencies. As shown in Fig. 4A, mutant mice spent significantly more time awake (53.8 ± 3.7%) compared to controls (32.6 ± 3.4%, *p* = 0.028), which was accompanied by a reduction in NREM sleep duration in mutants (41.7 ± 3.6%) relative to controls (59.4 ± 3.5%, *p* = 0.028). The proportion of REM sleep remained unchanged between groups. In addition, sleep onset latencies were altered in mutant mice (Fig. 4B). Indeed, the latency to REM sleep onset was significantly prolonged in mutants (206.3 ± 45.7 min) compared to controls (66.3 ± 17.9 min, *p* = 0.028, Fig. 4B). The latency to NREM sleep onset also showed a trend toward an increase (133.0 ± 36.8 min in mutants vs. 38.9 ± 16.3 min in controls, *p* = 0.057, Fig. 4B). These results indicate that mutant mice experience prolonged wakefulness and delayed sleep onset, suggesting disruptions in the neural mechanisms governing sleep initiation. To further investigate sleep microarchitecture, we analyzed the mean number and duration of wake, NREM, and REM sleep episodes. No significant differences were detected between groups (Supplementary Fig. 3A, B), suggesting that while the overall distribution of vigilance states is altered in mutant mice, sleep fragmentation remains unchanged.

**Fig. 4.**
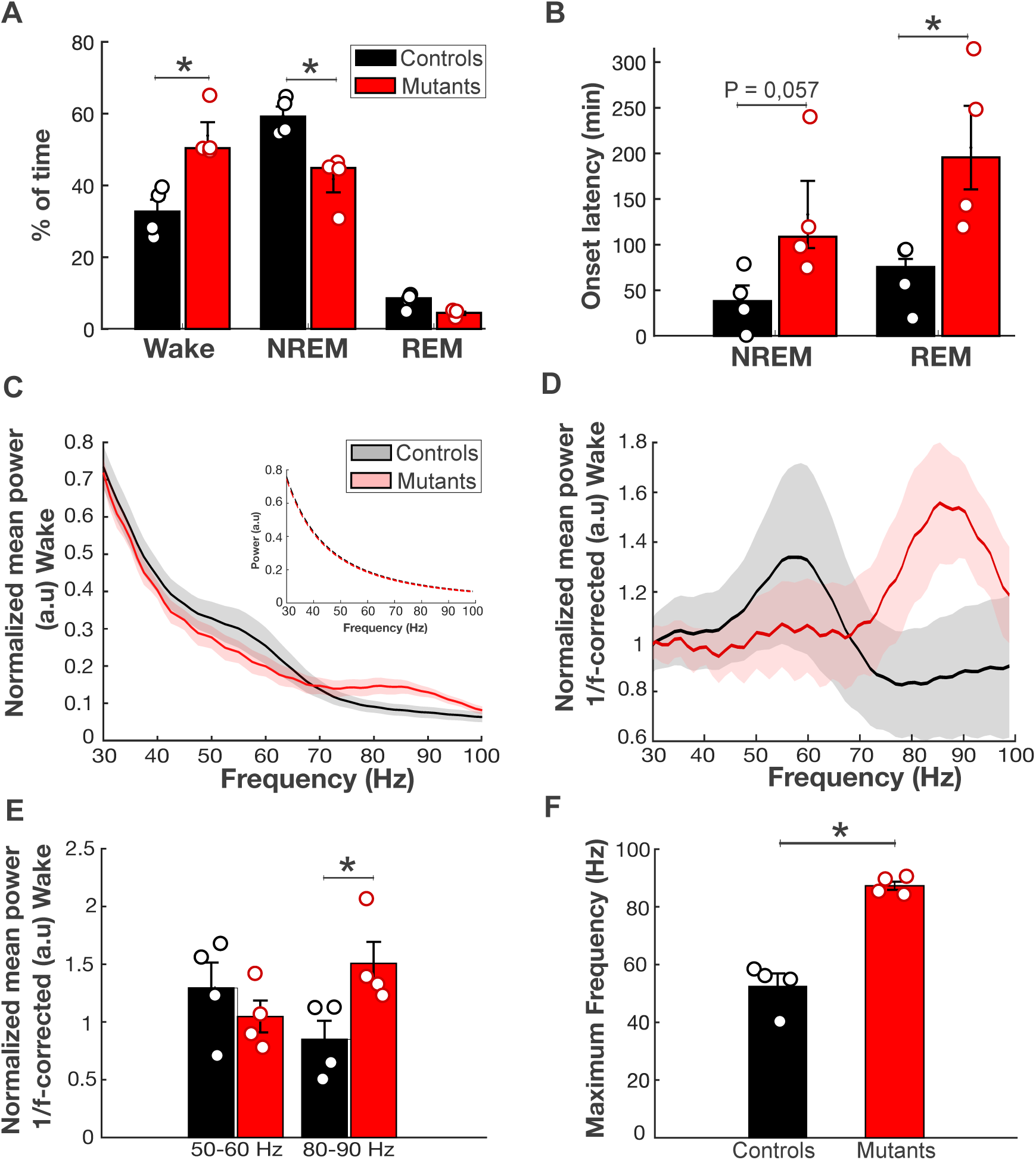
Altered diurnal sleep and gamma oscillatory activity in mutant compared to control mice. **A.** Percentage of diurnal time spent in wakefulness, NREM sleep, and REM sleep in control (black) and mutant (red) mice. Mutant mice exhibit significantly decreased NREM sleep and increased wake time compared to controls. **B.** Latency to the first NREM and REM sleep episodes. Mutant mice show a significant delay in REM sleep onset and a trend toward increased NREM sleep latency. **C.** Normalized mean power spectra during wakefulness for control and mutant mice. Shaded areas indicate the standard error of the mean (SEM). The inset shows the fitted 1/f curve used to correct for global power decay. **D.** Power spectra after 1/f correction, isolating oscillatory components from the broad 1/f decay. While control mice exhibit a peak in the 50–60 Hz gamma range, mutant mice display reduced lower gamma power and enhanced 80–90 Hz peak, suggesting altered gamma synchronization. **E.** Quantification of 1/f-corrected gamma power in the 50–60 Hz and 80–90 Hz ranges. Mutant mice exhibit a significant decrease in lower gamma power (50–60 Hz) and a significant increase in higher gamma power (80–90 Hz). **F.** Peak gamma frequency (40–100 Hz) for each group. The dominant gamma frequency is significantly higher in mutant mice compared to controls, suggesting a shift toward higher gamma frequencies in mutant mice. Bars represent mean ± SEM, and individual data points correspond to each mouse. Statistical comparisons were performed using the Mann-Whitney U test, and asterisks (*) indicate significant differences (*p* < 0.05) between groups (n = 4 per group).

To compare the spectral properties of control and mutant mice during wakefulness, we first computed the normalized mean power spectra for each group. Both control and mutant mice exhibit a characteristic 1/f power decay as frequency increases (Fig. 4C), which is usual in the brain. Mutant mice show a notable increase in power at the highest gamma frequencies compared to controls (Fig. 4C). To isolate genuine oscillatory components from the overall power decay, we applied a 1/f fitting procedure to the normalized mean spectra (Fig. 4C, inset). We then applied a 1/f correction to remove the influence of broadband power decay, allowing us to focus specifically on deviations from the expected 1/f distribution, which reflect true oscillatory activity rather than global shifts in power (Fig. 4D). After normalization, clear differences in gamma power emerge between groups, particularly in the 80–90 Hz range, where mutant mice exhibit a significantly higher power peak compared to controls (*p* = 0.03, Fig. 4E). Gamma power also tended to be lower in mutants within the 50–60 Hz range (Fig. 4D, E), but this difference was not statistically significant. Interestingly, the peak frequency of gamma oscillations appeared to be shifted toward higher frequencies in mutant mice compared to controls (Fig. 4D). To quantify this shift, we analyzed the dominant frequency of spectral power within the 40– 100 Hz range for each mouse. As shown in Fig. 4F, the peak gamma frequency was significantly higher in mutant mice (87.2 ± 1.4 Hz) compared to control mice (52.5 ± 4.1 Hz, *p* = 0.029). These results indicate that mutant mice exhibit a shift in gamma oscillatory dynamics, characterized by an increase in high-frequency gamma power (80–90 Hz) and a higher peak gamma frequency, suggesting potential alterations in inhibitory network activity.

To assess potential alterations in low-frequency oscillatory activity in mutant mice, we analyzed the normalized mean power spectra during wakefulness, NREM sleep, and REM sleep, focusing on two key frequency bands: delta (1–4 Hz) and theta (6–10 Hz). The spectral profiles of control and mutant mice exhibited a similar pattern across vigilance states (Supplementary Fig. 4). During wakefulness and REM sleep, both groups showed a dominant power peak in the theta band (Supplementary Fig. 4A, B, E, F), and quantitative analysis confirmed that delta and theta power did not differ significantly between groups. Similarly, during NREM sleep, both control and mutant mice displayed the expected high-amplitude delta activity associated with slow-wave sleep (Supplementary Fig. 4C), with no significant differences in delta or theta power between groups (Supplementary Fig. 4D). These findings indicate that low-frequency oscillations, including delta and theta rhythms, remain unaltered in mutant mice, suggesting that the observed electrophysiological differences are specific to gamma activity rather than broad changes in neocortical oscillatory dynamics.

### Behavioral consequences of TRα1^L400R^ expression in PV neurons

The results reported above show that gene expression, extracellular matrix glycosylation and electrophysiological properties are significantly altered in the neocortical PV neurons of *Thra^AMI/+^ Pvalb-Cre* mice, which might result in an alteration of the excitation-inhibition balance in the neocortex. We thus explored the possible behavioral consequences of *TRα1^L400R^* expression in PV neurons. Two groups of adult females, *Thra^AMI^*^/+^ *Pvalb-Cre* mice and *Thra^+/+^ Pvalb-Cre* control littermates, underwent a series of behavioral tests (Supplementary Table 3).

The most salient observation was that *Thra^AMI^*^/+^ *Pvalb-Cre* mice exhibited a higher level of general activity. They traveled significantly higher distances than control mice in the open-field, novel object recognition as well as social interaction tests (Fig. 5A, Supplementary Table 3). Moreover, in the acquisition phase of the test for novel object recognition, they spent significantly more time exploring objects (Fig. 5D). Finally, in the marble burying test, they buried significantly more marbles than controls (Fig. 5E). This reflects a higher level of digging behavior, which is a typical behavior that mice display when they are provided with sufficient substrate to displace (22).

**Fig. 5.**
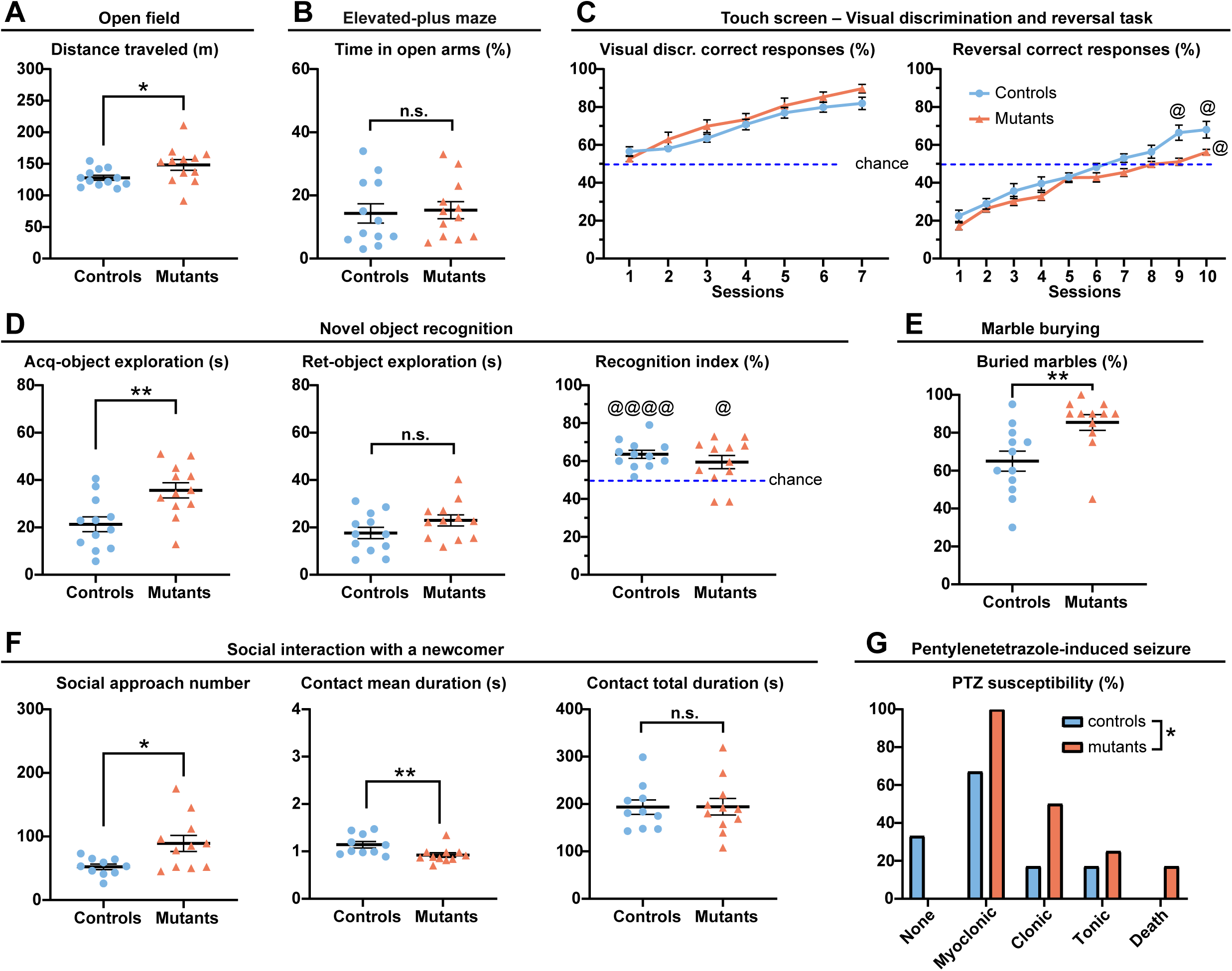
Behavioral alterations of mice expressing TRα1^L400R^ in PV neurons. **A.** Total distance traveled (m) in the open-field test was longer in mutant (*Thra^AMI^*^/*+*^ *Pvalb-Cre*) mice (n = 12) than that in control (*Thra^+^*^/*+*^ *Pvalb-Cre*) littermates (n = 12, unpaired two-tailed T-test). **B.** The time spent in open arms in the elevated-plus-maze test did not differ between controls (n = 12) and mutants (n = 12, unpaired two-tailed T-test). **C.** Touch-screen test: percentage of correct responses of controls (n = 11) and mutants (n = 11) in a visual discrimination task and in a reversal task (one sample T-test, @ *p* < 0.05 vs chance). **D.** Novel object recognition test: Exploration time (s) of the object during the acquisition (Acq-) and retention (Ret-) sessions, and recognition index (%) (controls n = 12, mutants n = 12, unpaired two-tailed T-test and one sample T-test, @ *p* < 0.05 vs chance and @@@@ *p* < 0.00001 vs chance). **E.** Marble-burying test: percentage of buried marbles (controls n = 12, mutants n = 12, Mann-Whitney test). **F.** Social interaction with a newcomer: Social approach number (unpaired two-tailed T-test), contact mean duration (Mann-Whitney test) and contact total duration (unpaired two-tailed T-test) (controls n = 10, mutants n = 11). **G.** Pentylenetetrazole-induced seizure test: percentage of mice showing myoclonic, clonic, tonic seizures and death (controls n = 12, mutants n = 12, Chi-square test). Data are presented as mean ± SEM and analyzed using the indicated tests (\**p* < 0.05, \*\**p* < 0.01). Also see Supplementary Table 3 for related analyses and datasets.

We next addressed the susceptibility of *Thra^AMI^*^/+^ *Pvalb-Cre* mice to an epileptogenic drug, the GABA receptor antagonist pentylenetetrazole (PTZ). In this test, they displayed more frequent and more severe seizures than controls (Fig. 5G).

Assessment of cognitive abilities revealed that, in spite of good learning abilities, *Thra^AMI^*^/+^ *Pvalb-Cre* mice exhibited less behavioral flexibility than controls. In the test for novel object recognition, control and mutant mice demonstrated a similar ability to discriminate a novel object from a known object (Fig. 5D). In a touch screen test, there was no difference between genotypes in visual discrimination learning (Fig. 5C). However, adapting to a change in the rule proved to be more difficult for mutant mice. In the reversal learning phase, mutant mice reached the learning criterion (% of correct responses higher than chance) during the 10^th^ session, whereas control mice reached the criterion during the 9^th^ session (Fig. 5C).

In a social interaction test, *Thra^AMI^*^/+^ *Pvalb-Cre* mice initiated significantly more social approaches and contacts (Fig. 5F, Supplementary Table 3). However, the mean duration of social contacts initiated by mutant mice was shorter, so that the total contact duration did not differ between genotypes. Thus, the higher numbers of social approaches and contacts in mutant mice appear to reflect their higher level of general activity rather than a higher degree of social motivation.

*Thra^AMI^*^/+^ *Pvalb-Cre* mice did not show any overt sign of increased anxiety. They spent the same amounts of time as controls in anxiogenic parts of test devices: open arms in the elevated-plus maze test (Fig. 5B), center of the arena in the open-field and social interaction tests (Supplementary Table 3).

## Discussion

Thyroid hormones play a crucial role in PV neuron maturation. Notably, TH/TRα1 signaling is instrumental in triggering the expression of the *Pvalb* gene (13). We demonstrate here that in the mouse neocortex, TH signaling is still required in PV neurons after they express the *Pvalb* gene, which is a late step in their maturation process. This expression starts in a protracted manner, at different time points in different neocortical areas, after the first postnatal week (7, 23). In the present study, we have found that blocking TH signaling in PV neurons at that stage, by expressing the dominant negative TRα1^L400R^ mutated receptor (*Thra^AMI^*^/+^ *Pvalb-Cre* mice), does not compromise the survival of PV neurons. It induces alterations in the expression of a small set of genes, impaired elaboration of PNNs with decreased PV neuron excitability and a change in cortical gamma oscillation dynamics. These network defects correlate with increased wakefulness, hyperactivity and increased susceptibility to epileptic seizure. The strength of the mouse model used in this study lies in its ability to specifically isolate the effects of disrupting TH signaling within PV neurons, ensuring that the observed phenotype stems directly from PV neuron dysfunction rather than from alterations in their microenvironment, which also plays a crucial role in regulating their physiology.

The mild consequences observed in mice expressing TRα1^L400R^ in PV neurons (*Thra^AMI^*^/+^ *Pvalb-Cre* mice, present paper) contrast with the postnatal lethality caused by spontaneous epileptic seizures observed in mice expressing TRα1^L400R^ in all GABAergic neurons from embryonic day 12.5 (13). A major difference between these two models is that in *Thra^AMI^*^/+^ *Pvalb-Cre* mice, the blockade of TH signaling is triggered at a late stage of PV neuron maturation, *i.e*. when *Pvalb* starts to be expressed, after the end of the first postnatal week. The different results obtained with *Thra^AMI^*^/+^ *Pvalb-Cre* and *Thra^AMI^*^/+^ *Gad2-Cre* mice might also reflect an indirect effect of TH/TRα1 signaling on PV neuron differentiation, via other GABAergic cell types. Indeed, the action of THs in PV neurons has been found to rely on a combination of cell-autonomous and non-cell-autonomous effects (24). However, compared to other cortical GABAergic neuron subtypes, PV neurons are, by far, the most sensitive to early TRα1^L400R^ expression in GABAergic neurons, as previously shown with histology and gene expression analyses (13). Therefore, the decreased severity of the phenotype observed in *Thra^AMI^*^/+^ *Pvalb-Cre* mice, compared to *Thra^AMI^*^/+^ *Gad2-Cre* mice, is likely to result from the delayed inhibition of TH signaling in PV neurons, rather than from its restriction to a fraction of the GABAergic neuronal population. To our knowledge, there is no indication that THs also directly regulate the differentiation of other neocortical GABAergic neurons in the neocortex. Thus, TH signaling does not only play a major role for PV neuron differentiation before the onset of *Pvalb* expression, but is also required for the lifelong maintenance of specific characteristics of PV neurons.

Gene expression analysis indicates that THs directly regulate the transcription of only a small set of genes in PV neurons, of which several influence the elaboration of the PNN. Among these genes, several contribute to PNN elaboration, such as *Acan, Sema7a* and *Npnt* genes, that encode extra-cellular matrix proteins: aggrecan is a major component of the PNN, semaphorin 7a is anchored to the cell membrane, and nephronectin is an integrin-interacting glycoprotein. Furthermore, the deregulation of *Galnt14* and *Has3* genes, respectively encoding N-acetylgalactosaminyl transferase and hyaluronan synthase, might account for alterations of the carbohydrate fraction of PNNs, revealed by WFA staining. However, mechanisms that are more indirect can also be at work. For example, down-regulation of *Me1, Ampd2* and *Nudt7*, which encode enzymes, suggests alterations of the energy metabolism in PV neurons, which might translate into PNN defects. Such a metabolic hypothesis has been previously put forward to explain the PNN phenotype of mice lacking the PGC1α TR coactivator in cortical GABAergic neurons (25). This PNN defect could explain why PV neurons expressing TRα1^L400R^ have a reduced sensitivity to stimulation, as revealed by patch–clamp analysis. However, additional molecular mechanisms might also be involved in the reduced excitability of PV neurons in *Thra^AMI^*^/+^ *Pvalb-Cre* mice, like the deregulated expression of the *Tesc* gene, which encodes tescalcin, a calcium-binding protein, and of the *Cacnb3* gene, encoding a voltage-dependent calcium channel subunit. Of note, *Cacnb3* was previously found to be up-regulated after TH treatment in PV neurons of the motor cortex (26) and it has emerged as a genetic risk factor for bipolar disorder in humans (27). As a whole, the changes in gene expression induced by the blockade of TH signaling in PV neurons are consistent with the observed decrease in PNN abundance and in PV neuron excitability.

The alteration of electrophysiological properties of PV neurons caused by TRα1^L400R^ expression was accompanied by an increase in high-frequency gamma power (80–90 Hz) in the SomCx. This observation is counterintuitive because PV neurons are known to drive gamma oscillations in the cortex (4). However, this apparent discrepancy can be explained by considering the broader network impact of PV interneuron dysfunction. Reduced excitability of PV interneurons decreases inhibitory control over pyramidal neurons, which is expected to cause a disinhibition and increased excitatory drive. This elevated excitatory activity likely enhances gamma oscillations, particularly at higher frequencies, consistent with findings from studies on neocortical circuit dynamics (4, 28). Furthermore, impaired PV interneuron function could disrupt the precise timing required for effective inhibition, altering network synchronization and shifting gamma oscillations toward higher peak frequencies, as observed in our mutant mice. In addition, other inhibitory neurons, notably somatostatin-expressing interneurons, also significantly modulate cortical oscillatory activity (29) and they might be recruited to compensate PV neurons defects. The shift toward higher gamma frequencies likely results from chronic PV interneuron dysfunction. As a whole, changes in cortical gamma oscillations are likely to impact cognitive processing (30).

*In vivo* recordings also show that, compared to controls, mutant mice expressing TRα1^L400R^ selectively in PV neurons exhibit significantly increased wakefulness during the day, along with a prolonged sleep latency. A possible causal relationship between hypothyroidism and short sleep duration has been suggested in humans, but the link could not be firmly established, as sleep quality and duration rely on a combination of factors (31). The fact that *Thra^AMI^*^/+^ *Pvalb-Cre* mice exhibit hyperactivity and increased wakefulness is consistent with the increase in high-frequency gamma power (80–90 Hz) and with the shift of the peak gamma frequency to higher values during wakefulness. Indeed, EEG activity recorded in human patients with insomnia have previously revealed that elevated activity in high-frequency bands typically reflects hypervigilance (32). Similarly, the increased high-frequency gamma activity observed in our mutant mice likely represents a comparable hyperarousal state, which might translate in heightened motor activity, as observed in behavior tests. This hyperarousal might compromise sleep initiation, thereby contributing to prolonged sleep latency and heightened wakefulness (33). As a whole, our results highlight that TH signaling in PV neurons plays a direct role in sleep-wake regulation. In addition, the changes in cortical gamma oscillations induced by blocking TH/TRα1 signaling in PV neurons are consistent with the hyperactivity and reduced sleep duration observed in mutant mice.

The behavioral phenotype of mice expressing TRα1^L400R^ only in PV neurons was mainly characterized by a general increase in activity, reduced behavioral flexibility and an increased susceptibility to pentylenetetrazole-induced seizures (Fig. 5). These findings likely derive from an imbalance between excitation and inhibition in the brain, and suggest that altered TH signaling in PV neurons underlies part of the symptoms experienced by patients with altered TH signaling. Of note, an involvement of cortical PV neuron activity in reward-related behavioral flexibility had been reported before (34). The behavioral defects observed in the present study differ notably from those reported for mice expressing TRα1^R384C^ in the whole body, which display high anxiety, memory defects (35, 36) and reduced seizure susceptibility (37). The susceptibility to epilepsy is an indication of an increase in the excitation drive, compared to the inhibition drive in neural circuits, a feature observed in many neuropsychiatric disorders (1, 38). Thus, the heightened susceptibility to epilepsy that we have observed is congruent with the reduced excitability of PV neurons in mice expressing TRα1^L400R^ only in PV neurons. Moreover, as THs also exert a cell-autonomous influence in other brain cells, such as adult excitatory cortical neurons (26) and glial cells (39), the behavioral traits observed in mice expressing TRα1^R384C^ in all cells reflect a combination of direct and indirect effects of the mutation, starting from prenatal developmental stages. As a consequence, the deregulation of gene expression in adult PV neurons is unlikely to play a major part in the phenotype of TRα1^R384C^ mice. By contrast, the present paper uncovers specific consequences induced by disrupting TH/TRα1 signaling in PV neurons, notably in terms of general activity, behavioral flexibility and susceptibility to epilepsy.

The genomic, histological, electrophysiological and behavioral traits induced in mice by blocking TH/TRα1 signaling in PV neurons form a coherent whole, suggesting that in the context of reduced TH signaling, PV interneuron dysfunction induces complex network adaptations in the neocortex. These observations provide new insights into the role of TH signaling in maintaining inhibitory network function, with potential implications for neurological and psychiatric disorders involving altered cortical network oscillations, such as autism-spectrum disorders, schizophrenia, Alzheimer’s disease, attention-deficit disorders or epilepsy. In humans, adult onset of hypothyroidism may cause memory impairment, depression (40), forgetfulness, difficulty focusing and “brain fog” (41). It is also linked to poor sleep quality (42). In these patients, concomitant fatigue, muscle weakness, bradycardia, and decrease in resting energy expenditure, can indirectly cause the cognitive and mood disorders. The current study highlights that the decrease in TH signaling in PV neurons alone accounts for at least some of the symptoms of adult hypothyroidism, which could result from an imbalance between neuronal excitation and inhibition.

### Study limitations

The identification of TR target genes in PV neurons is incomplete, because we were not able to perform a Chip-Seq analysis to firmly establish the positions of TR binding sites in the genome, due to an insufficient amount of chromatin extracted from sorted PV neuron nuclei.

For technical reasons, whole-cell current-clamp recording was carried out in juvenile mice, so it cannot be excluded that PV neuron excitability might evolve at later stages.

The present study was focused on the neocortex, but as parvalbumin-expressing neurons are present in other brain areas, it is likely that blocking TH/TRα1 signaling in PV neurons also affected other neural circuits. As a consequence, part of the behavioral phenotype observed in *Thra^AMI^*^/+^ *Pvalb-Cre* mice might originate from alterations in brain areas other than the neocortex.

Further investigations are needed to establish links between the observed changes in gene expression caused by TRα1^L400R^ expression in the neocortex and PV neuron functional defects.

## Methods

### Mouse models and treatments

Experiments involving the use of live animals for the current project were approved by local ethics committees (C2EA015 and C2EA017) and subsequently authorized by the French Ministry of Research (Projects #33279 and #38581). Mice were bred and maintained at Plateau de Biologie Expérimentale de la Souris (SFR BioSciences Gerland - Lyon Sud, France). Unless otherwise specified, mice of both sexes were used in the experiments.

The *Thra^AMI^* allele allows the expression of the dominant-negative TRα1^L400R^ after the Cre/loxP-mediated deletion of a cassette with polyadenylation signals (14). The *ROSA-tdTomato^lox^*reporter transgene (also known as Ai9, MGI Cat# 4436851, RRID: MGI: 4436851) drives the Cre-dependent expression of a cytoplasmic red fluorescent protein (43) . The *ROSA-GS-GFP-L10^lox^* transgene encodes the GS-EGFP-L10a protein, which is a green fluorescent protein fused to the N-terminus part of the large subunit ribosomal protein L10a, mainly localized in cell nuclei. This transgene allows for a strong and specific staining of the nuclei (44). The additional N-terminal GS tag, combines a fragment of protein G and a fragment of streptavidin suitable for protein purification (45). The entire reading frame and a drug-selection gene (NeoR) were inserted in the pHL-HH vector construct (46) which contains 2 LoxP sequences and 2 LoxP^2272^ sequences (44). The resulting vector was inserted into the ROSA26 locus of mouse embryonic stem cells by homologous recombination (Supplementary Fig. 1A). The combination of LoxP and LoxP^2272^ sites is organized so that Cre/LoxP recombination, taking place only between identical sequences, results in a two-step inversion/deletion (Supplementary Fig. 1B). This design makes the expression of GS-EGFP-L10a strictly dependent on Cre recombination (46). All transgenes were in C57Bl6/J genetic background. *Pvalb-Cre* is a knock-in allele in which the Cre recombinase reading frame is inserted into the *Pvalb* gene in order to restrict the expression of the recombinase to PV neurons (47). The *ROSA-tdTomato^lox^* reporter system has allowed us to assess the specificity of PV neuron targeting. Of note, we have observed that in *ROSA-tdTomato^lox/+^ Pvalb-Cre* mice, Cre-mediated recombination occurred not only in PV neurons throughout the brain, but also in cerebellar granule cells.

Hypothyroidism was induced in adult *ROSA-GS-GFP-L10^lox/+^ Pvalb-Cre* mice by feeding them for 2 weeks with iodine deficient food supplemented with 0.15% of propylthiouracyl (PTU, Envigo ref TD.95125). TH levels were restored by a single intraperitoneal injection in half of the mice (40 μg T4 + 4 μg T3 dissolved in 200 μL of phosphate buffer saline, all chemicals from Sigma Aldrich France). The other PTU-treated mice were injected with 200 µL of phosphate-buffered saline.

### Nuclei sorting

Individual neocortex was frozen in liquid nitrogen immediately after dissection. Samples were thawed and resuspended in 1.5 mL of lysis buffer (Nuclei EZ Lysis Buffer, Sigma, ref N3408) and transferred in a 2 mL Dounce tissue grinder (Sigma, ref D8938) for cell lysis (25 strokes with A pestle and 25 strokes with B pestle, on ice). Suspensions were transferred in tubes containing 2.5 mL of cold lysis buffer and centrifuged (5 min 500 g 4°C, swinging rotor). Supernatants were discarded and nuclei pellets were washed once with 4 mL Nuclei EZ Lysis Buffer and once with 4 mL of Nuclei Suspension Buffer (NSB) (PBS w/o Mg and Ca; 0.01% BSA; Biolabs ref B9000S; 40 U/mL RNAsin; Promega, ref N251B). For each of these washes, pellets were first resuspended by pipetting 10 times with 1 mL of buffer, after which the remaining 3 mL of buffer were added. Final pellets were resuspended in 0.5 mL NSB containing DAPI (1 µL at 1 mg/mL) and were filtrated in tubes with 35 μm cell strainer (Corning, ref 352235). For each condition, Fluorescence-Activated Cell Sorting (FACS Aria IIµ cytometer (BD)) was used to isolate all GFP positive nuclei. These were sorted in low binding Eppendorf tubes containing 0.5 mL NSB and centrifuged (5 min 20 000 g). Supernatants were discarded and nuclei pellets were resuspended in 350 μL RNA lysis buffer containing β-mercaptoethanol for RNA extraction (Qiagen RNeasy Microkit, ref 74004).

### RNA-Seq analysis

Nuclear RNA was extracted from sorted nuclei (RNeasy Micro Kit, Qiagen ref 74004) and quantified using Tapestation4150 (Agilent). Then, 1 ng of RNA was reverse-transcribed using the SMART-Seq v4 Low Input RNA kit (Takara). cDNA was quantified and qualified using Qubit (Invitrogen) and Tapestation4150 (Agilent). Libraries were then prepared from 1 ng of cDNA using the Nextera XT DNA Library Kit (Illumina). Libraries were sequenced (> 2.10^7^ reads/library) on a Nextseq500 DNA sequencer (Illumina). Reads were aligned on the mouse genome (mm10 GRCm38 release) with Bowtie2 (Galaxy Version 2.4.2 + galaxy 0) (48). Count table was prepared using htseq-count (Galaxy Version 0.9.1 + galaxy 1, mode union, feature type: gene) (49). Differential gene expression analysis was performed with DEseq2 (Galaxy Version 2.1.8.3) (50) using the following thresholds: False Discovery Rate < 0.05; *p*-adjusted value < 0.05; expression > 10 reads per million; fold-change > 1.5). Hierarchical clustering was performed using Euclidian distance and Ward’s distance with Clustvis (51).

### ATAC-seq analysis

Neocortex nuclei were purified as above and collected by centrifugation (500 g 5 min) for immediate use in ATAC Library preparation (Tagmentase-loaded; Diagenode). The nuclei pellets were suspended in 25 μL 2xTagmentation Buffer (Diagenode) before addition of 22.5 μL of H_2_0 and 2.5 μL of Tn5 transposase. This mix was incubated under agitation (1400 rpm) for 30 min at 37°C. DNA fragments were purified using the MinElute PCR Purification Kit (Qiagen). The eluate (10 µL) was amplified by PCR after the addition of 12.5 µL H_2_0, 25 µL NEBNext® High-Fidelity 2xPCR Master Mix (New England Biolabs) and 2.5 µL of index primers (Nextera® XT Index kit v2; Illumina) as follows: 72°C for 5 min, 98°C for 30 s, followed by 5 cycles (98°C 10 s, 63°C 30 s, 72°C for 60 s). A preliminary qPCR was then performed with a fraction of the reaction product to determine the optimal number of amplification cycles needed for a second round of amplification without saturation: to this end 5 µL of iQ SYBR Green Supermix (Biorad) were added to 5 µL preamplified DNA and used for qPCR (98°C 30 s denaturation), followed by 20 cycles of amplification: (98°C 10 s, 63°C 30 s, and 72°C 60 s). The entire preamplified DNA was then amplified (usually 9–11 cycles). DNA from the final reaction was purified using 1.8x SPRI magnetic beads (Beckman Coulter) quantified with Qubit 4 fluorometer (Invitrogen) and qualified using Tapestation4150 (Agilent). Libraries were mixed in equimolar amounts and sequenced (> 2.10^8^ reads/library; Nextseq500 DNA sequencer Illumina). Reads were aligned on the mouse genome (mm10 GRCm38 release) with Bowtie2 (Galaxy Version 2.4.2 + galaxy 0) (48). Peaks were identified using MACS2 Galaxy Version 2.2.9.1 + galaxy 0). MACS2 was also used for differential occupancy analysis using a set of reference library to define background.

### Brain slice collection

Each mouse was given a lethal intraperitoneal injection (6 mL/kg) of a mixture of ketamine (33 mg/mL) and xylazine (6.7 mg/mL). The thorax was opened and each mouse was perfused with 3.7% paraformaldehyde in 0.1 M phosphate buffer at room temperature. Each brain was dissected out, immersed in fixative at 4°C for 3 h and then in phosphate buffered saline (PBS) at 4°C until sectioning. Coronal sections (50 μm thick) were cut with the aid of a vibrating microtome (Integraslice 7550 SPDS, Campden Instruments, Loughborough, UK), in PBS at room temperature. Brain sections were stored at -20°C in cryoprotectant (30% ethylene glycol and 20% glycerol in 10 mM low-salt PBS).

### Brain slice histology

Immunohistochemistry was performed on free-floating brain sections. For PV labeling, a mouse anti-PV primary antibody (PARV19, Sigma P3088, 1:2000), and a secondary antibody made in donkey (anti-mouse DyLight 633, ThermoFisher Scientific, 1:1000) were used. For the concurrent PNN labeling, a biotinylated Wisteria floribunda lectin (WFL/WFA, Vector Laboratories B-1355, 20 µg/mL), and Streptavidin coupled to DyLight 488 (Vector Laboratories, SA-5488, 1:1000) were used. For ROS labeling, a mouse monoclonal 8-Hydroxy-2’-deoxyguanosine primary antibody (8-oxo-dG, Abcam ab48508, 1:2000), and a secondary antibody made in donkey (anti-mouse DyLight 488, ThermoFisher Scientific, 1:1000) were used. Non-specific binding sites were blocked by incubating sections for 1 h in PBS with 1% BSA and 0.2% Triton X100. Brain sections were incubated overnight at 4°C with PV antibody and biotinylated WFA, which were diluted in PBS with 1% BSA, 0.2% Triton X-100 and 1% dimethyl sulfoxide. Sections were washed in PBS and further incubated for 15 min at room temperature with DAPI (4’, 6-diamidino-2-phenylindole, 1:5000, Sigma). Incubation with the secondary antibody lasted for 3 h at room temperature. Sections were mounted in Fluoroshield^TM^ (Sigma), coverslipped and imaged using an inverted confocal microscope (Zeiss LSM 780 and Zeiss LSM 800).

### Image analysis

Image analysis was performed using ImageJ. Final images resulted from the z projection of 3 optical sections (Maximum intensity tool) that were 2 µm apart (the resulting images thus reflected the fluorescence collected over a thickness of 4 µm). Tomato+, PV+ and WFA+ cells were counted with the Cell Counter in ImageJ plugin. For the 8-oxo-DG labeling experiment, Tomato+ 8-oxo-DG+ cells were detected from final images using the Cellpose wrapper for Fiji (52). Numbers of double-labelled cells per slice, 8-oxo-DG labeling intensity in Tomato+ cells and 8-oxo-DG labeling intensity outside Tomato+ cells were measured automatically from 3-4 slices per animal. The code of the original macro may be found here: https://github.com/jbrocardplatim/Macro_8oxo.

### Acute slice preparation

Juvenile *Thra^AMI/+^ Pvalb-Cre RosatdTomato* and *Pvalb-Cre RosatdTomato* littermates (16-24-day-old) were deeply anesthetized with isoflurane and decapitated. The brains were quickly removed and placed into cold oxygenated artificial CerebroSpinal Fluid (aCSF): 126 mM NaCl, 2.5 mM KCl, 1.25 mM NaH_2_PO_4_, 2 mM CaCl_2_, 1 mM MgCl_2_, 26 mM NaHCO_3_, 10 mM glucose, 15 mM sucrose, and 1 mM kynurenic acid (nonspecific glutamate receptor antagonist; Sigma). Coronal slices (300 μm thick) from mouse SomCx were prepared as described previously (20). Slices were cut with a vibratome (VT1000S; Leica), transferred to a holding chamber containing aCSF saturated with O_2_/CO_2_ (95%/5%), and held at room temperature.

### Whole-cell electrophysiological recording

Individual slices were transferred to a submerged recording chamber and perfused (1–2 ml/min) with oxygenated aCSF. Patch pipettes (∼10 MΩ) pulled from borosilicate glass were filled with 12 µl of internal solution: 144 mM K-gluconate, 3 mM MgCl_2_, 0.5 mM EGTA, 10 mM HEPES, pH 7.2 (285/295 mOsm). Neurons were first identified in the slice by their tdTomato fluorescence and then patch under infrared differential interference contrast illumination (SliceScope Pro, Scientifica). Whole-cell recordings in current-clamp mode were performed at room temperature (24.5 ± 1.5°C) using a patch-clamp amplifier (MultiClamp 700B; Molecular Devices). Data were filtered at 10 kHz and digitized at 100 kHz using an acquisition board (Digidata 1440A; Molecular Devices) attached to a personal computer running pClamp 10.7 software package (Molecular Devices). All membrane potentials were corrected for theoretical liquid junction potential (-15.6 mV).

### Stereotaxic surgery for in vivo extracellular local field potential recording

8-week-old male *Thra^AMI^*^/+^ *Pvalb-Cre* mice and *Thra^+/+^ Pvalb-Cre* control littermates were anesthetized in an induction chamber with 3% isoflurane. Following induction, they received analgesia via a subcutaneous injection of buprenorphine (0.1 mg/kg) and were placed in a stereotaxic frame (David Kopf Instruments, USA). Throughout the procedure, anesthesia was maintained with 1.5–2% isoflurane, and core body temperature was regulated at 37°C using a heating pad. Prior to the skin incision, a local subcutaneous injection of lidocaine (4–5 mg/kg) was administered for additional analgesia. Tungsten wire electrodes were bilaterally implanted in the olfactory bulb (OB) (AP: +4.3, ML: ±0.5, DV: −1.5), the CA1 hippocampal layer (AP: −2.2, ML: ±2.0, DV: −1.0), and the SomCx (AP: −2.0, ML: ±3.5, DV: −1.2). A reference electrode was positioned on the cerebellar surface. Additionally, two wires were inserted into the nuchal muscles to record the electromyogram (EMG). Upon completion of the implantation, electrodes were secured to the skull using wax and dental cement (SuperBond C & B). The electrode leads were then connected to an EIB32 interface board, which was fixed onto the skull using Parafilm and dental cement. Brain activity recordings commenced after a minimum recovery period of one week.

### In vivo Electrophysiological recordings

Neural signals from all implanted electrodes were recorded using an Intan Technologies amplifier chip (RHD2216) at a sampling rate of 20 kHz. Local field potentials (LFPs) were downsampled and stored at 1,250 Hz for further analysis. Signal processing and data analysis were conducted using custom-made MATLAB scripts, based on publicly available toolboxes, including <u>Battaglia’s computing tools</u> and the <u>FMA Toolbox</u>. LFPs were recorded using tungsten wire electrodes with PFA insulation (0.002” bare diameter).

### Sleep scoring

Automatic vigilance state classification was performed using a MATLAB-based algorithm developed in-house (53) This approach integrates two parallel data streams: LFP recordings from the hippocampus and accelerometer measurements from the head-mounted EIB sensor, which finely monitor movement of the animal along the x, y, and z axes. Wakefulness versus sleep is distinguished based on accelerometer activity levels, with sustained low activity indicating sleep. The differentiation between NREM and REM sleep was based on the theta (5–10 Hz) / delta (2–4 Hz) power ratio in the hippocampus. Specifically, NREM sleep is characterized by a predominance of slow waves, reflected by high delta power, whereas REM sleep is marked by strong theta oscillations.

### Behavior tests

Adult female *Thra^AMI^*^/+^ *Pvalb-Cre* mice and *Thra^+/+^ Pvalb-Cre* control littermates were transferred to PHENOMIN-ICS platform (Illkirch, France), where they underwent the following series of behavioral tests (Supplementary Table 3). 1. Open field: Each mouse was placed in the periphery of a 44*44 cm arena (Panlab, Barcelona, Spain), under 150 Lux illumination, and allowed to explore freely the apparatus for 30 min. The distance traveled, the number of rearings as well as the time spent in the central and peripheral regions were recorded (periphery width = 8 cm). 2. Elevated-plus maze: The apparatus comprised two open arms (30 × 5 cm) opposite one to the other and crossed by two closed arms (30 x 5 x 15 cm), all arms being equipped with infrared captors allowing automatic detection of the position of the mouse (Imetronic, Pessac, France). Mice were individually tested for 5 min. The following parameters were recorded: number of entries into and time spent in each arm, head dips, rearings, stretched-attend posture. 3. Visual discrimination and reversal task: Mice were tested in Mouse Touch Screen Systems (Campden Instruments). In order to increase the motivation of the animals, the drinking water in the housing cages was acidified during test days. After pre-training to learn to touch an image that appeared on the screen in order to obtain a reward, mice were tested in a visual discrimination task. Touching the correct image was rewarded with strawberry milk (12% sucrose). Touching the incorrect image was punished with a timeout. Once the task was learnt, the correct image became incorrect and vice versa. Each animal performed a session of 1 hour maximum per day, involving up to 60 trials per session, during which the percentage of correct responses was recorded. 4. Novel object recognition: Mice were first habituated for 10 min in a closed circular arena (30 cm diameter and 50 cm height, homogeneously illuminated at 20 Lux). The next day, mice were submitted to a 10-min acquisition trial, during which they were placed in the same arena in presence of two identical objects (A). The time spent exploring the objects (sniffing) was manually recorded. Two hours later, mice were placed again in the same arena, where one of the objects had been replaced by a novel object (B). During this 10-min retention trial, the times spent exploring the objects (tA and tB) were recorded. A recognition index (RI) was defined as (tB / (tA + tB)) x100. Locomotor activity was recorded with EthoVision XT video tracking system (Noldus, Wageningen, Netherlands). 5. Marble burying: Mice were tested individually for 15 minutes in a cage (Tecniplast GM500, illuminated at 25 lux) containing 3 cm high clean bedding and 20 black glass marbles (10 mm diameter) arranged in 5 lines of 4 marbles. EthoVision XT video tracking system (Noldus, Wageningen, Netherlands) was used for video recording. The number of marbles fully covered by bedding was measured every minute, between the 5th and 15th minutes of the test. 6. Social interaction with a newcomer: Mice were tested in an open arena of 50*50 cm with sawdust. The animals were previously equipped with an RFID chip so that each animal could be tracked independently and its social behaviors recorded by the Live Mouse Tracker system (54). For the habituation session, the mouse under test was allowed to freely explore the arena for 20 minutes. An unknown female adult C57Bl/6 mouse (“newcomer”) was then placed in the arena for 30 min. The social behaviors, including social approaches, contacts, following the other mouse, were recorded as well as the distance traveled by each mouse. 7. Pentylenetetrazole-induced seizure: Each mouse received an intraperitoneal injection of pentylenetetrazole (with 40 mg/kg) and was placed in a translucent cage. Different types of seizures were manually recorded for 20 minutes following the injection: myoclonic (short jerking in parts of the body), clonic (periods of shaking or jerking parts on the body), tonic (muscles in the body become stiff).

### Statistical analyses

Data are expressed as mean ± SEM. Data analyses were performed with Excel 2016 and GraphPad Prism software (v.8.). The level of significance was set at *p* < 0.05. For histological analysis, the densities of Tomato+, PV+, WFA+, 8-oxo-DG+ cells, and their co-localizations were counted in the whole area of one image taken in the SomCx area. 3–4 images in both hemispheres were analyzed for each mouse. Quantitative data were analyzed using two-tailed unpaired *T-*test. In electrophysiological analysis, 37 parameters were manually defined as described (20), adopting Petilla terminology (21). The normality was analyzed using Shapiro-Wilk test. Quantitative data were analyzed using Mann-Whitney test or two-tailed unpaired *T-*test. For mouse behavior analysis, the normality was analyzed using Shapiro-Wilk test. Quantitative data were analyzed using two-tailed unpaired *T*-test or Mann-Whitney test and two-way ANOVA followed by Sidak’s multiple comparisons test. For the comparison with chance, one group *T*-test was used. Outliers were identified with the ROUT method (Q = 1%). A Chi-square test was used to analyze the susceptibility to pentylenetetrazole-induced seizures. The analysis of in vivo local field potential recordings was performed in the MATLAB environment (MathWorks). This included the classification of vigilance states and the computation of spectral power in the SomCx. For statistical comparisons, non-parametric analyses were conducted using the Mann-Whitney U test to assess differences across vigilance states and spectral power distributions in the SomCx.

## Acknowledgements

We acknowledge the contribution of SFR Biosciences (Université Claude Bernard Lyon 1, CNRS UAR3444, Inserm US8, ENS de Lyon): Nadine Aguilera and the Plateau de Biologie Expérimentale de la Souris (ANIRA-PBES) for mouse breeding; ANIRA-CYTOMETRIE and particularly Sébastien Dussurgey for nuclei sorting; ANIRA-AGC for mouse genotyping. We acknowledge Benjamin Gillet and Sandrine Hughes of the deep sequencing facility (PSI IGFL, Lyon). We thank Hugues Jacobs, Olivia Wendling, Emel Laghouati, Tania Sorg-Guss and Yann Hérault for the behavioral tests performed at Mouse Clinical Institute (Strasbourg, France). We acknowledge the contributions of the CELPHEDIA Infrastructure (http://www.celphedia.eu/), especially the center AniRA in Lyon. We also thank Isabelle Dusart (Institut de Biologie Paris Seine, France) for helpful discussions. We sincerely thank Karim Benchenane, Sophie Bagur, and Baptiste Maheo (Brain Plasticity Unit, eMOBBS team) for their valuable advice and discussions regarding the analysis of in vivo electrophysiological data. We also thank Ke Sun for writing Python programs (https://jupyter.org/try-jupyter/lab/index.html) that automated part of the data analysis.

We acknowledge Chinese Scholarship Council for a scholarship to J.R. (# 202106140005) to do her PhD project in ENS de Lyon (https://www.chinesescholarshipcouncil.com/fr/). F.F received research fundings from European Union’s Horizon 2020 research and innovation program (https://research-andinnovation.ec.europa.eu/funding/funding-opportunities/funding-programmes-and-opencalls/horizon-2020_en) under grant agreement No. 825753 (ERGO), Agence Nationale de la Recherche (https://anr.fr/, Thyromut2 program; ANR-15-CE14-0011-01), Joint Research Institute for Science and Society (JORISS 2022, https://www.enslyon.fr/indexation/structures-affiliees-lens-et-partenaires/joint-research-institutescience-and-society) and the PHENOMIN program of AAP AIN INSB CNRS 2022 (https://www.insb.cnrs.fr/fr/appel-projets-developpement-des-infrastructuresnationales-de-recherche).

## Author contribution

Conceptualization: F.F., S.R., J.R., B.C., T.G., F.R. Investigation: J.R., R.G., D.A., A.D., T.G, J.B., S.M., D.L., F.R, F.F., S.R. Writing-Original Draft: J.R., F.F, S.R. Writing – Review & Editing: All authors. Funding Acquisition: F.F., J.W. Supervision: S.R., F.F., D.L., B.C., T.G.

