## Supplementary Fig. 1 for "Thyroid hormones maintain parvalbumin neuron functions in the mouse neocortex"

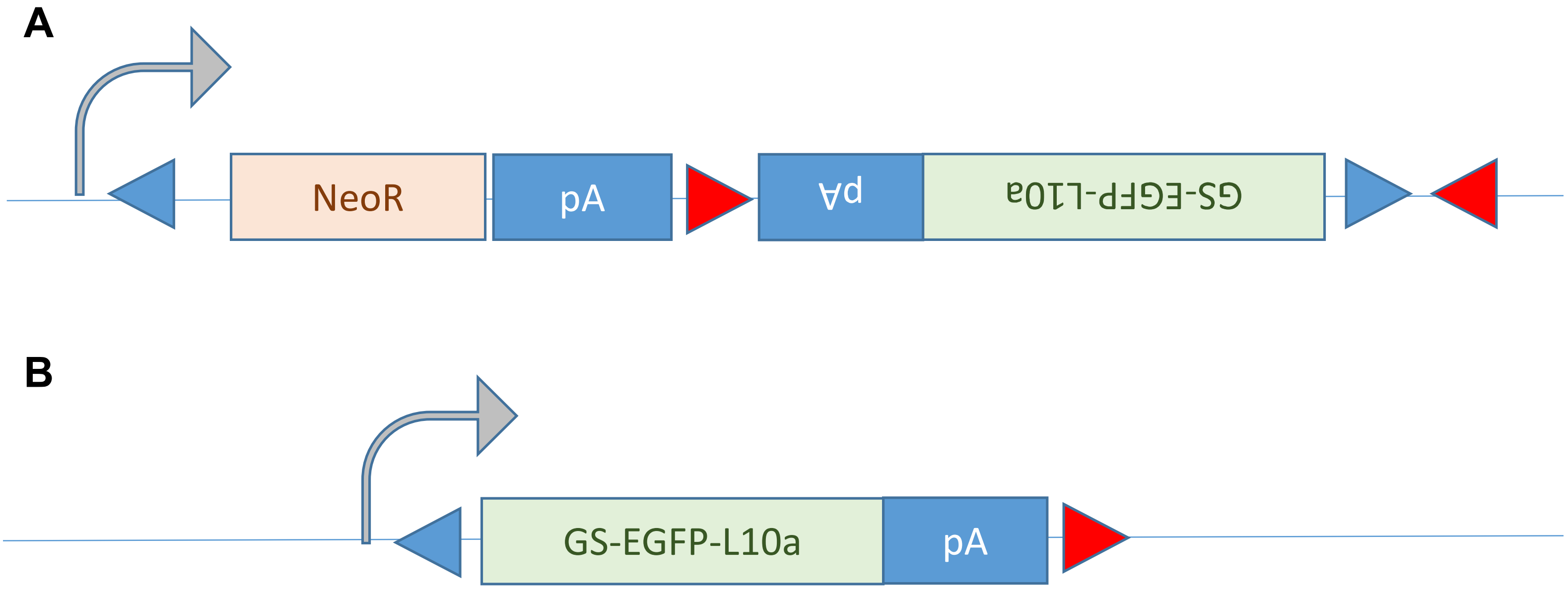


**Supplementary Fig. 1 | *ROSA-GSGFPL10*$^{lox}$ reporter construct that was used to sort PV neuron nuclei in *Thra*$^{AMI}$ *ROSA -GSGFPL10*$^{lox}$ *Pvalb-Cre* mice. A** EGFP, a green fluorescent protein, was fused to the N-terminus part of the large ribosomal protein subunit L10a, allowing a strong and specific staining of nuclei (Heiman et al. 2008). An additional N-terminal GS tag, which combines a fragment of G protein and a fragment of streptavidin (Burckstummer et al., 2006), was added for subsequent protein purification. The entire reading frame was inserted in the *pHL-HH* vector (Fehling et al., 2003). This construct contains a drug-selection gene (*NeoR*), two loxP sequences (blue triangles) and two loxP$^{2272}$ sequences (red triangles). This vector was inserted into the *ROSA26* locus of mouse embryonic stem cells by homologous recombination. Before Cre-mediated recombination, the *ROSA* promoter drives the expression of the *NeoR* drug resistance gene. **B** After a two-step recombination mediated by the Cre recombinase between LoxP (blue) and loxP$^{2272}$ (red) sequences, the fragment encoding *NeoR* is deleted and the fragment encoding the GS-EGFP-L10a fusion protein is inverted. This results in the expression of a fluorescent protein (GS-EGFP-L10a), which is predominantly nuclear and strictly dependent on Cre-mediated recombination (Fehling et al., 2003).
