## Supplementary Fig. 2 for "Thyroid hormones maintain parvalbumin neuron functions in the mouse neocortex"

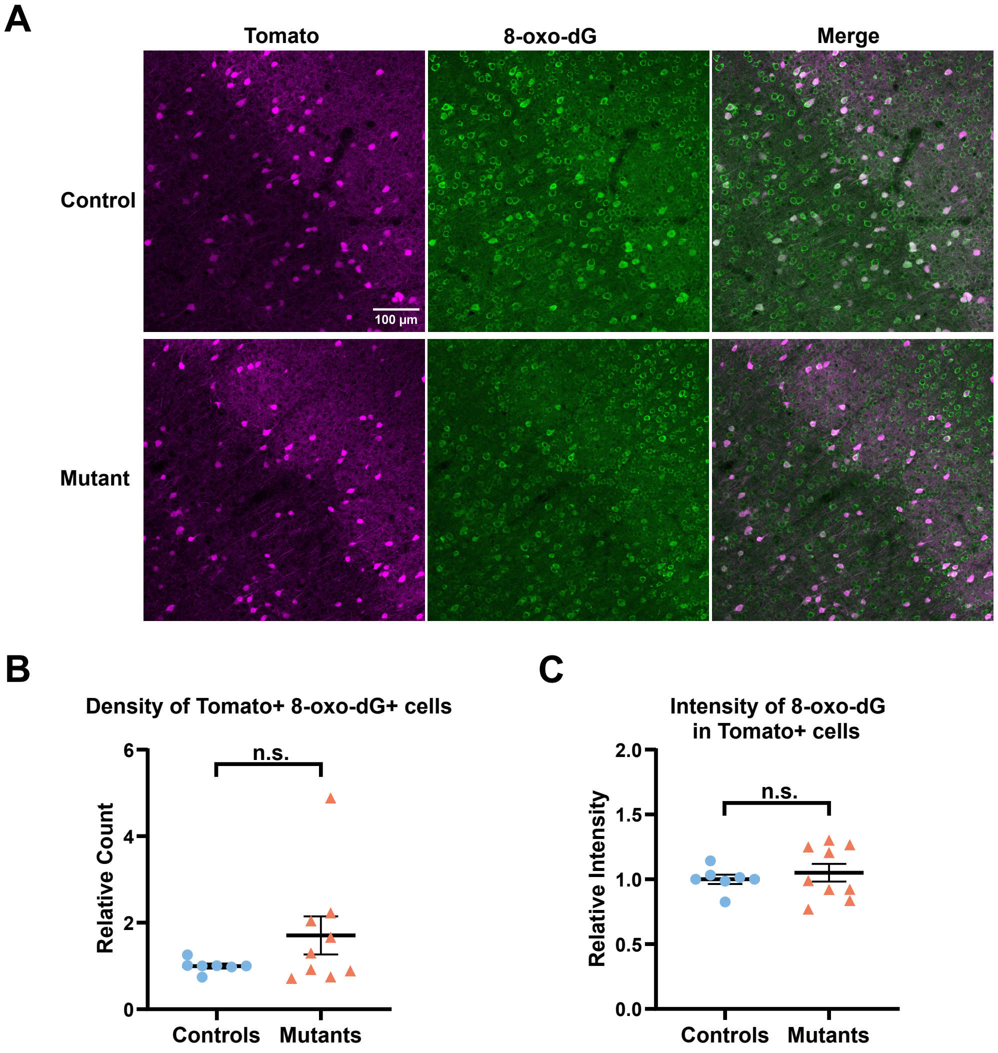


**Supplementary Fig. 2 | Investigation of oxidative stress levels in mice expressing TRα1$^{L400R}$ in PV neurons. A** Immunohistochemistry of the oxidative stress marker 8-oxo-dG in PV neurons of the somatosensory cortex in *Thra$^{AMI/+}$ ROSA-tdTomato$^{lox/+}$ Pvalb-Cre* mice and *Thra$^{+/+}$ ROSA-tdTomato$^{lox/+}$ Pvalb-Cre* control littermates. **B, C** The relative density of Tomato+ 8-oxo-dG+ cells and the relative intensity of 8-oxo-dG in Tomato+ cells (relative to the intensity of the signalin the rest of the section) did not differ between the two groups. Data are presented as mean ± SEM and were analyzed using unpaired two-tailed T-test.

In view of the absence of difference between the two groups, we sought to assess the power of our analysis, focusing on the relative intensity of 8-oxo-dG labelling in Tomato+ cells, compared to that in the rest of the section. To assess the minimum detectable difference between control and mutant groups, the 95% confidence interval (CI) for the mean difference may be calculated. With mean values of 1.000 ± 0.093 (mean ± SD) for the control group (n = 7) and 1.051 ± 0.205 for the mutant group (n = 9), the mean difference was 0.051 with a 95% CI: -0.129 to 0.231. Since this confidence interval includes zero, it confirms that there is no statistically significant difference between the groups; the width of this confidence interval indicates that the study had sufficient power to detect a difference greater than 0.231.
