## Supplementary Fig. 3 for "Thyroid hormones maintain parvalbumin neuron functions in the mouse neocortex"

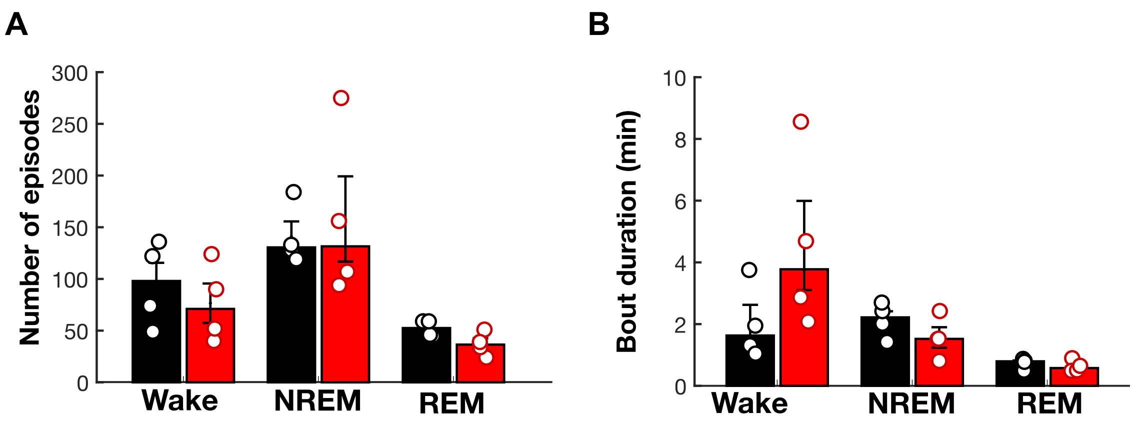


**Supplementary Fig. 3 | Analysis of sleep-wake episode frequency and duration in mutant and control mice. A** Mean number of episodes for each vigilance state (wake, NREM, and REM). **B** Mean bout duration of each state. Bars represent mean ± SEM for control (black) and mutant (red) mice (n = 4 per group).
