## Supplementary Fig. 4 for "Thyroid hormones maintain parvalbumin neuron functions in the mouse neocortex"

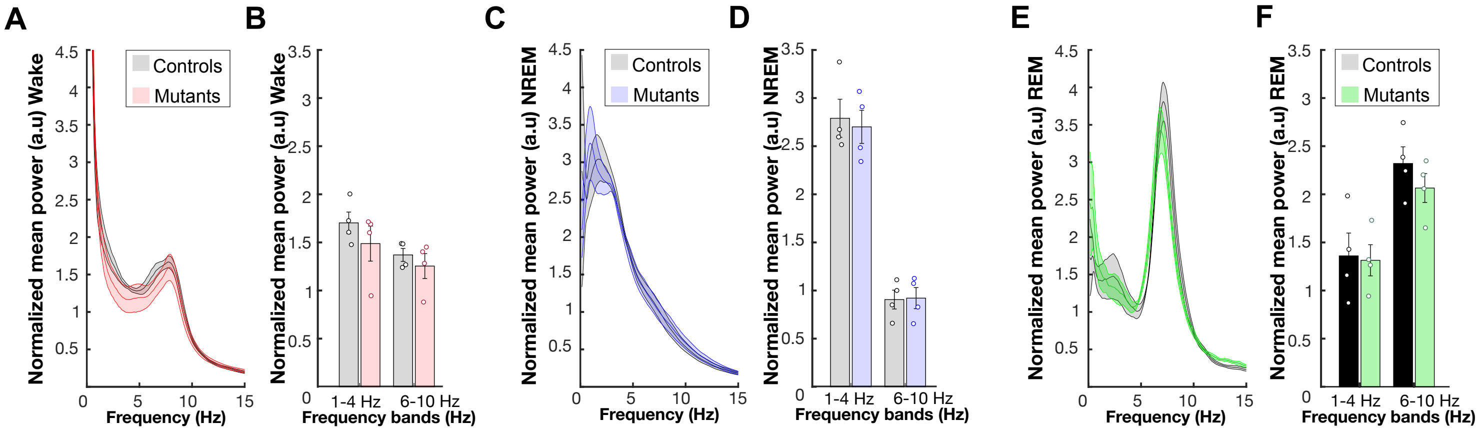


**Supplementary Fig. 4 | Delta and Theta power remain unaltered across vigilance states in mutant mice. A, C, E** Averaged power spectra (0 – 15 Hz) in the somatosensory cortex during wakefulness (A), NREM sleep (C), and REM sleep (E). Gray curves represent control mice, while colored curves represent mutant mice (blue for NREM, green for REM). Both groups display similar spectral profiles across vigilance states, with dominant theta activity during wakefulness and REM sleep and delta activity during NREM sleep. **B, D, F** Quantification of delta (1 – 4 Hz) and theta (6 – 10 Hz) power for each vigilance state. Bars represent mean ± SEM for control (gray) and mutant (colored) mice (blue for NREM, green for REM). Statistical comparisons using the Mann-Whitney U test revealed no significant differences between groups (n = 4 per group).
