## Supplementary Table 2 for "Thyroid hormones maintain parvalbumin neuron functions in the mouse neocortex"

**Supplementary Table 2. Statistical result for 26 essential electrophysiological parameters.** Control n = 19 (cells from 8 mice), Mutant n = 35 (cells from 10 mice).

| **Phase** | **Parameters** | **mean ± SEM** | | ***p* value** | |
| --- | --- | --- | --- | --- | --- |
|  |  | **Control** | **Mutant** | **T-test** | **Mann-Whitney** |
| Subthreshold properties | RMP (mV) | -78.68 ± 1.83 | -80.70 ± 0.94 |  | 0.72 |
|  | R_m_ (MΩ) | 269.20 ± 33.08 | 202.69 ± 16.18 |  | 0.12 |
|  | t_IC_ (ms) | 15.32 ± 2.40 | 12.61 ± 0.84 |  | 0.29 |
|  | C_m_ (pF) | 63.47 ± 7.45 | 68.06 ± 3.94 |  | 0.20 |
|  | Δ G_sag_ (%) | 0.08 ± 0.02 | 0.09 ± 0.01 |  | 0.51 |
| Near threshold firing properties | **Rheobase (pA)** | **102.71 ± 14.57** | **153.66 ± 14.05** | **0.02 *** |  |
|  | 1^st^ spike delay (ms) | 277.42 ± 58.54 | 183.11 ± 35.87 |  | 0.20 |
|  | m_threshold_ (Hz/s) | -18.97 ± 12.52 | -26.89 ± 10.70 |  | 0.83 |
|  | C_threshold_ (Hz) | 23.21 ± 3.80 | 29.44 ± 3.91 |  | 0.38 |
|  | Hump (mV) | 0.86 ± 0.15 | 0.63 ± 0.06 |  | 0.30 |
| Saturating firing properties | A_sat_ (Hz) | 34.10 ± 2.15 | 28.16 ± 3.23 | 0.21 |  |
|  | t_sat_ (ms) | 26.84 ± 7.37 | 59.38 ± 21.26 |  | 0.92 |
|  | C_sat_ (Hz) | 170.02 ± 10.60 | 165.80 ± 9.17 | 0.78 |  |
|  | m_sat_ (Hz/s) | -28.81 ± 3.45 | -32.63 ± 4.16 |  | 0.98 |
| Action Potential Waveform | A1 (mV) | 54.83 ± 1.49 | 51.72 ± 1.65 | 0.22 |  |
|  | AHP1f (mV) | -26.01 ± 0.68 | -26.48 ± 0.64 | 0.64 |  |
|  | T_AHP1f_ (ms) | 2.63 ± 0.19 | 2.38 ± 0.14 | 0.27 |  |
|  | D1 (ms) | 0.61 ± 0.03 | 0.57 ± 0.03 |  | 0.14 |
|  | **Threshold AP1 (mV)** | **-35.45 ± 0.97** | **-32.28 ± 0.94** | **0.04 *** |  |
|  | A2 (mV) | 53.23 ± 1.28 | 50.58 ± 1.44 | 0.23 |  |
|  | AHP2f (mV) | -26.48 ± 0.65 | -27.23 ± 0.56 | 0.41 |  |
|  | T_AHP2f_ (ms) | 2.62 ± 0.16 | 2.45 ± 0.15 |  | 0.30 |
|  | D2 (ms) | 0.62 ± 0.03 | 0.57 ± 0.03 |  | 0.13 |
|  | **Threshold AP2 (mV)** | **-34.52 ± 0.90** | **-31.51 ± 0.84** | **0.03 *** |  |
|  | Amp, Red, | 0.03 ± 0.01 | 0.02 ± 0.01 | 0.58 |  |
|  | Dur, Inc, | 0.02 ± 0.01 | 0.01 ± 0.00 |  | 0.46 |
