## Supplementary Table 3 for "Thyroid hormones maintain parvalbumin neuron functions in the mouse neocortex"

**Supplementary Table 3. Statistical results for key parameters in behavioral tests.**

| **Age (week)** | **Test** | **Parameters** | **Control** | | **Mutant** | | ***p* value** | |
| --- | --- | --- | --- | --- | --- | --- | --- | --- |
|  |  |  | **N** | **Mean ± SEM** | **N** | **Mean ± SEM** | **T-test** | **Mann-Whitney** |
| 17 | Open Field | Distance traveled (m) | 12 | 127.95 ± 10.49 | 12 | 148.40 ± 8.54 | **0.04 *** |  |
|  |  | Number of rearings | 12 | 219.25 ± 15.71 | 12 | 270.33 ± 21.64 | 0.07 |  |
|  |  | % Time in center | 12 | 21.46 ± 1.30 | 12 | 24.95 ± 2.02 |  | 0.36 |
|  |  | Entries in center | 12 | 198.08 ± 7.03 | 12 | 241.42 ± 12.55 | **0.006 **** |  |
|  |  | Distance in center (m) | 12 | 43.88 ± 3.80 | 12 | 54.75 ± 3.88 | **0.02 *** |  |
|  |  | Distance in periphery (m) | 12 | 84.07 ± 7.23 | 12 | 93.65 ± 6.87 | 0.23 |  |
| 17 | Elevated Plus Maze | Open Arm Entries | 12 | 4.67 ± 0.63 | 12 | 5.08 ± 0.65 | 0.65 |  |
|  |  | Closed Arm Entries | 12 | 11.92 ± 0.84 | 12 | 12.92 ± 0.87 | 0.42 |  |
|  |  | Total Arm Entries | 12 | 16.58 ± 1.12 | 12 | 18.00 ± 1.27 | 0.41 |  |
|  |  | % Open Entries | 12 | 27.42 ± 2.84 | 12 | 27.67 ± 2.49 | 0.95 |  |
|  |  | Attempts | 12 | 20.75 ± 2.12 | 12 | 25.67 ± 1.44 | 0.07 |  |
|  |  | Head-Dips | 12 | 6.92 ± 1.12 | 12 | 5.00 ± 0.99 | 0.21 |  |
|  |  | Stretched-Attend Posture | 12 | 6.08 ± 0.72 | 12 | 6.33 ± 0.72 | 0.81 |  |
|  |  | Open Arm Entry Latency (s) | 12 | 25.78 ± 9.29 | 12 | 40.05 ± 20.43 |  | 0.88 |
|  |  | Open Time (s) | 12 | 43.08 ± 9.19 | 12 | 46.17 ± 8.01 | 0.80 |  |
|  |  | Closed Time (s) | 12 | 154.83 ± 5.66 | 12 | 147.75 ± 9.91 | 0.54 |  |
|  |  | Center Time (s) | 12 | 102.50 ± 8.87 | 12 | 106.75 ± 7.92 | 0.72 |  |
|  |  | % Time in Open Arms | 12 | 14.33 ± 3.04 | 12 | 15.33 ± 2.68 | 0.81 |  |
|  |  | % Open Entries / (Open + Closed) | 12 | 20.42 ± 3.83 | 12 | 23.67 ± 3.86 | 0.56 |  |
|  |  | Open Arm Time (proximal) | 12 | 19.50 ± 2.93 | 12 | 21.08 ± 2.27 |  | 0.60 |
|  |  | Open Arm Time (distal) | 12 | 23.42 ± 7.25 | 12 | 24.92 ± 7.33 |  | > 0.99 |
| 18-22 | Touch Screen | Nb of trials before criterion for Visual Discrimination | 10 | 162 ± 15.62 | 12 | 130 ± 19.31 |  | 0.19 |
|  |  | Nb of trials before criterion for Reversal Task | 11 | 517.36 ± 24.10 | 11 | 583.73 ± 8.16 |  | **0.03 *** |
| 23 | Novel Object Recognition | Habituation Distance Traveled (m) | 12 | 35.70 ± 1.32 | 12 | 45.35 ± 3.93 | **0.03 *** |  |
|  |  | Acquisition Distance Traveled (m) | 12 | 23.72 ± 1.28 | 12 | 29.76 ± 2.45 |  | **0.04 *** |
|  |  | Retention Distance Traveled (m) | 12 | 18.67 ± 0.99 | 12 | 23.33 ± 2.53 |  | **0.03 *** |
| 24 | Marble Burying | % Buried marbles (15 min) | 12 | 65.00 ± 5.33 | 12 | 85.42 ± 4.15 |  | **0.005 **** |
| **Age (week)** | **Test** | **Parameters** | **Control** | | **Mutant** | | ***p* value** | |
|  |  |  | **N** | **Mean ± SEM** | **N** | **Mean ± SEM** | **T-test** | **Mann-Whitney** |
| 25 | Social Interaction | Habituation Total Distance (m) | 10 | 77.60 ± 2.76 | 11 | 95.33 ± 8.24 |  | **0.04 *** |
|  |  | Habituation Center Distance (m) | 10 | 13.16 ± 0.96 | 11 | 19.14 ± 3.17 |  | 0.0845 |
|  |  | Habituation % Distance in Center | 10 | 16.78 ± 0.79 | 11 | 19.25 ± 1.59 | 0.1946 |  |
|  |  | Habituation Center Time (s) | 10 | 97.36 ± 5.92 | 11 | 122.25 ± 13.74 | 0.1251 |  |
|  |  | Habituation % Time in Center | 10 | 8.11 ± 0.49 | 11 | 10.19 ± 1.15 | 0.1251 |  |
|  |  | Distance traveled (m) | 10 | 99.24 ± 4.13 | 10 | 133.30 ± 13.53 | **0.003 **** |  |
|  |  | Social Approach Nb | 10 | 52.30 ± 4.37 | 11 | 89.00 ± 12.59 | **0.02 *** |  |
|  |  | Social Approach Mean Duration (s) | 10 | 0.70 ± 0.03 | 11 | 0.73 ± 0.04 | 0.55 |  |
|  |  | Social Approach Total Duration (s) | 10 | 36.72 ± 3.51 | 11 | 66.84 ± 11.90 |  | 0.05 |
|  |  | Contact Nb | 10 | 170.20 ± 9.66 | 11 | 210.55 ± 15.46 | **0.04 *** |  |
|  |  | Contact Mean Duration (s) | 10 | 1.14 ± 0.07 | 11 | 0.92 ± 0.05 |  | **0.006 **** |
|  |  | Contact Total Duration (s) | 10 | 193.35 ± 15.29 | 11 | 194.32 ± 17.60 | 0.97 |  |
|  |  | Oral-Genital Nb | 10 | 78.10 ± 7.61 | 11 | 97.91 ± 8.83 | 0.11 |  |
|  |  | Oral-Genital Mean Duration (s) | 10 | 0.39 ± 0.01 | 11 | 0.37 ± 0.02 | 0.39 |  |
|  |  | Oral-Genital Total Duration (s) | 10 | 30.01 ± 2.70 | 11 | 35.72 ± 3.43 | 0.21 |  |
|  |  | Oral-Oral Nb | 10 | 83.60 ± 7.58 | 11 | 86.09 ± 7.50 | 0.82 |  |
|  |  | Oral-Oral Mean Duration (s) | 10 | 0.45 ± 0.02 | 11 | 0.43 ± 0.02 | 0.27 |  |
|  |  | Oral-Oral Total Duration (s) | 10 | 37.45 ± 2.96 | 11 | 35.97 ± 2.50 | 0.71 |  |
|  |  | Side by Side Nb | 10 | 44.60 ± 6.37 | 11 | 51.00 ± 9.89 |  | 0.80 |
|  |  | Side by Side Mean Duration (s) | 10 | 0.42 ± 0.02 | 11 | 0.38 ± 0.01 | 0.06 |  |
|  |  | Side by Side Total Duration (s) | 10 | 18.77 ± 2.75 | 11 | 19.13 ± 3.53 | 0.94 |  |
|  |  | Follow Nb | 10 | 2.80 ± 0.85 | 11 | 5.18 ± 1.27 | 0.14 |  |
|  |  | Follow Mean Duration (s) | 10 | 0.40 ± 0.07 | 11 | 0.41 ± 0.05 | 0.95 |  |
|  |  | Follow Total Duration (s) | 10 | 1.34 ± 0.37 | 11 | 2.45 ± 0.68 | 0.18 |  |
| 33 | Pentylenetetrazole induced Seizure | PTZ susceptibility | 12 |  | 12 |  | **0.05 *** | Chi-square |
